# A novel antidepressant acting *via* allosteric inhibition of GluN2D-incorporated NMDA receptors at GABAergic interneurons

**DOI:** 10.1101/2022.11.06.514872

**Authors:** Jilin Zhang, Jinjin Duan, Luyu Ye, Wei Li, Haitao Zhou, Fang Liu, Tian Xiaoting, Yang Xie, Yiming Huang, Yidi Sun, Hu Zhou, Chenggang Huang, Yang Li, Shujia Zhu, Fei Guo

## Abstract

N-methyl-D-aspartate receptors (NMDARs) are glutamate-gated calcium-permeable excitatory channels. They have attracted great interest as potential targets for the treatment of depression in recent years. NMDARs typically assemble as heterotetramers composed of two GluN1 and two alternative GluN2 (2A-2D) subunits, the latter of which endow various subtypes with diverse gating and pharmacological properties. To date, limited molecules with GluN2 specificity have been identified to show antidepressant effects. Here, we identify a compound termed YY-23 extracted from *Rhizoma Anemarrhenae* allosterically inhibited the channel activities of GluN2C- or GluN2D-incorporated NMDARs, an effect that was presumably influenced by the S2 segment in the ligand-binding domain of the GluN2 subunit. We found that prefrontal GluN2D-containing NMDARs were predominantly expressed at GABAergic interneurons rather than pyramidal neurons. Furthermore, we revealed that YY-23 suppressed the activity of GluN2D-containing NMDARs and GABAergic neurotransmission in the medial prefrontal cortex (mPFC). As a consequence, this GABAergic disinhibition facilitated the excitatory transmission. Behavioural experiments showed that YY-23 acted as a rapid antidepressant in both stress-naïve and stressed animal models, which was abolished in *Grin2d-knockout* mice. Together, our findings suggest that GluN2D-incorporated NMDARs on GABAergic interneurons might be promising therapeutic targets for the treatment of depression.

## Introduction

Major depressive disorder (MDD) is one of the most common psychiatric diseases (1). After decades of preclinical and clinical research, the neurobiological basis and aetiological mechanism of MDD remain elusive, probably due to the heterogeneous genetic and environmental correlates, the lack of potential biomarkers, and the scarcity of representative animal models. Most antidepressants available in the clinic that have been developed based on the classic monoaminergic theory, however, produce low remission rates with long response lag times, moderate pharmacotherapeutic effectiveness and high individual heterogeneity (2). Thus, there is an urgent need for the development of faster-acting and more efficacious therapeutic agents to treat patients with MDD, especially those who are refractory to conventional antidepressants.

In recent years, a wealth of compelling studies have implied that impairment of the glutamatergic system is likely to underlie the pathophysiology of depression (3, 4). This mounting evidence has mainly been obtained from studies on the rapid-acting antidepressant ketamine, which is an NMDAR channel blocker (5–8). It has been suggested that ketamine treatment enhanced glutamate transmission in the prefrontal cortex (PFC) and hippocampus in rodents and humans through the blockade of NMDA receptors (NMDARs) on GABAergic interneurons (9, 10). This process instantly triggers dendritic spine regrowth (11) and cortical excitability (6, 7). Thus, NMDAR antagonism has received much attention as a theory for the development of rapid antidepressants.

NMDARs are glycine- and glutamate-coactivated ion channels that participate in synaptic neurotransmission and plasticity in the vertebrate brain (12). Typical NMDARs are tetramers composed of two glycine-bound GluN1 and two alternative glutamate-bound GluN2 (2A-2D) subunits. The four GluN2 subunits display distinct temporal and spatial expression patterns and contribute to various biophysical and pharmacological properties of NMDARs (13, 14). Notably, cell-type-specific NMDAR subtypes have been approved as molecular targets for the antidepressant action of ketamine and rapastinel; the latter is a positive allosteric modulator of NMDARs (15).

Our previous study identified a compound named YY-23 that selectively inhibited the activity of NMDARs (not AMPARs) in cultured hippocampal neurons (16, 17). In this study, we aimed to assess whether YY-23-mediated inhibition of NMDARs displays subtype selectivity and to understand the molecular mechanism underlying its fast antidepressant effects *in vivo*. Interestingly, we found that YY-23 selectively inhibited the gating activity of GluN2C- or GluN2D-containing NMDARs in both voltage- and agonist concentration-independent manners, acting as an allosteric inhibitor. Electrophysiological recordings together with transgenic animal behaviour studies further revealed that the GluN2D subunit at GABAergic interneurons in the medial prefrontal cortex (mPFC) played a pivotal role in mediating the antidepressant effects of YY-23.

## Materials and methods

All plasmids encoding wild-type NMDAR genes were generously provided by Dr. Pierre Paoletti (Institut de Biologie de l’Ecole Normale Supérieure, France). Other plasmids encoding chimeric, truncated and mutant NMDARs were constructed by Gibson assembly (NEBuilder® M5520AA) or site-directed mutagenesis. HPLC–MS/MS was used to measure YY-23 characterization. A Lance Ultra cAMP Kit and FLIPR® Calcium 4 Assay Kit (Molecule Devices) were used to screen pharmacological targets for YY-23 among monoaminergic receptors. For mRNA expression assessment in the mPFC, fluorescent oligonucleotide probes against *Grin2d, Slc32a1*, and *Slc17a7* mRNAs were used to perform multiplex fluorescence in situ hybridization. Two-electrode voltage-clamp recording and patch-clamp recording were performed as previously reported (18). Whole-cell patch-clamp recordings were performed on neurons from prelimbic mPFC slices to assay NMDA-induced currents, action potentials (APs), and excitatory postsynaptic currents (EPSCs). Field excitatory postsynaptic potentials (fEPSPs) in the mPFC were determined by placing the stimulating electrode on prelimbic mPFC layer VI and placing the recording electrode on layer II/III. MaxQuant (http://maxquant.org/, version 1.6.5.0) was used to analyse the MS/MS spectral data yielded by nanoflow liquid chromatography tandem mass spectrometry. All animal studies and experimental procedures were approved by the Animal Care Committees of the Shanghai Institute of Materia Medica and the Center for Excellence in Brain Science and Intelligence Technology, Chinese Academy of Sciences. *Grin2d-knockout* mice were generated by CRISPR/CAS9 based on C57BL/6N mice. To assess depression-related behaviours, the forced swimming test (FST), tail suspension test (TST), open field test (OFT), novelty-suppressed feeding test (NSFT) and social interaction (SI) test were performed on stress-naïve or stressed animals. All the details of the methods are described in the Supplementary Methods and Materials.

## Results

### YY-23 is an allosteric inhibitor of GluN2C- or GluN2D-containing NMDARs

As described in our previous report (17, 19), YY-23 was obtained by acid hydrolysis of timosaponin B3 or chemical synthesis from timosaponin B2 (Fig. S1 and Table S1). YY-23 was isolated as a white amorphous powder, [α]D −66.0 (c 0.05, pyridine), with a molecular formula of C_33_H_54_O_8_ as determined by positive-ion HR-ESI-TOF-MS (detected m/z: 601.3713 [M+Na]+; Calcd. For C_33_H_54_O_8_Na: 601.3711). The ^13^C-NMR spectrum and distortionless enhancement by polarization transfer (DEPT) experiment showed the presence of 4 methyl groups, 12 methylene groups, 13 methine groups, and 4 quaternary carbons. The presence of a β-glycopyranosyl moiety was determined by the anomeric proton at δ_H_ 4.87 (J = 7.7 Hz) and the carbon signal at δ_C_ 105.19. The NMR spectral data of YY-23 were very similar to those of timosaponin BIII, indicating that the two have the same basic skeleton. In the HMBC spectra, long-range correlations were observed from δ_H_ 4.87 (H-1’) to δ_C_ 75.22 (C-26), and *vice versa*. Accordingly, the structure of YY-23 was elucidated and named (25S)-3,26-dihydroxy-20(22)-ene-5β-furost-26-β-D-glucopyranoside (Fig. 1a).

**Figure 1.**
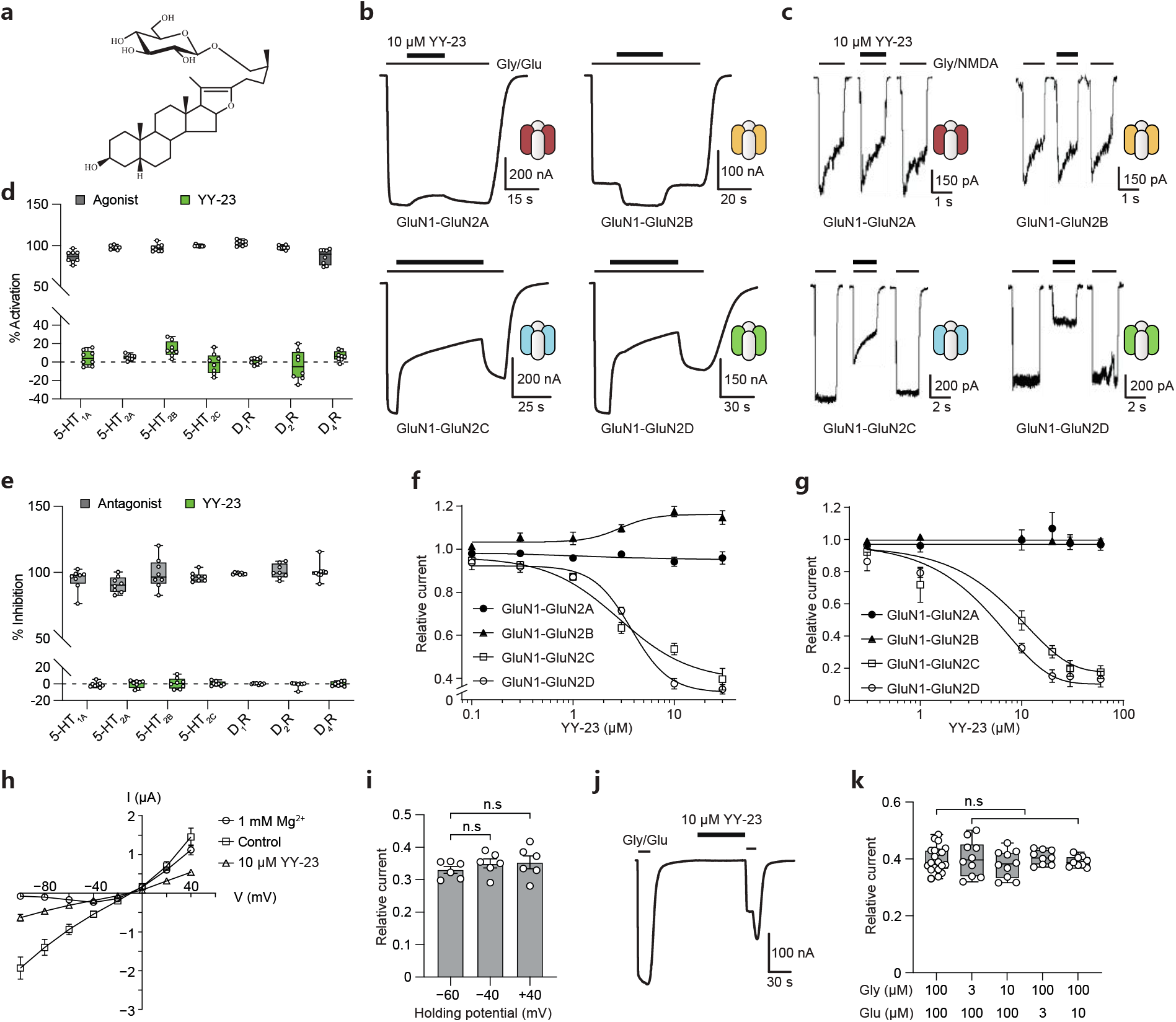
Subunit selectivity of YY-23 action. (a) Chemical structure of YY-23. (b and c) Representative current traces of recombinant NMDARs expressed in Xenopus oocytes recorded by TEVC during coapplication of 100 μM glutamate, 100 μM glycine and 10 μM YY-23 and in HEK293T cells recorded by the whole-cell voltage-clamp method during coapplication of 20 μM glycine, 500 μM NMDA and 10 μM YY-23. (d and e) A FLIPR® Calcium 4 Assay kit was used to analyse the possible activating (d) and inhibitory (e) effects of YY-23 on 5HT_1A_, 5HT_2B_ and 5HT_2_C. A Lance Ultra cAMP kit was used to test the effects of compounds on 5HT_1A_, D_1_R, D_2_R and D_4_R in agonist (d) and (e) antagonist mode. Data with normal and abnormal distributions were analysed by unpaired t test and Mann–Whitney test, respectively. Concentration–response curves of the effects of YY-23 on recombinant NMDARs expressed in Xenopus oocytes (f, data are shown as the mean ± SD) and HEK293T cells (g, data are shown as the mean ± SEM). The responses of saturating agonists were normalized to 1. (h) Current-voltage (I-V) relationships of GluN1-GluN2D receptors in the absence (control, square) and presence of 10 μM YY-23 (triangle) or 1 mM Mg^2+^ (circle), upon saturating glycine and glutamate (100 μM each). (i) Statistical analysis of YY-23-mediated inhibition at holding potentials of −60, −40 and +40 mV. The data were analysed by one-way ANOVA. (j) Recording trace for application of 10 μM YY-23 to GluN1-GluN2D receptors in the absence of agonists, with continuous application of YY-23 for 1 min. (k) Inhibition of GluN1-GluN2D receptor activity by YY-23 with different concentrations of glycine or glutamate. The currents induced by different agonist solutions in the absence of YY-23 were normalized to 1. The data were analysed by Brown-Forsythe ANOVA test. Each point represents a single cell. The bar graphs show the mean ± SEM. In the box plot graphs, the bars show the min/max values, the boxes show the lower/upper quartiles, and the lines represent the median values. n.s means no significance (p > 0.05).

First, we screened whether this compound targets monoaminergic receptors. FLIPR® Calcium 4 Assay Kits were used to screen the 5HT_2A_, 5HT_2B_ and 5HT_2C_ serotonin receptors, and Lance Ultra cAMP Kits were used to screen the 5HT_1A_, D1R, D2R and D4R dopamine receptors. Application of 10 μM YY-23 resulted in no significant activation or inhibition of these receptors (Fig. 1d, e and Table. S2). This indicated that the monoaminergic receptors were unlikely to be the targets of YY-23. Based on our previous data, YY-23 selectively inhibits NMDA- but not AMPA-induced currents in cultured hippocampal neurons (17). We next investigated whether YY-23-mediated inhibition displays subtype specificity by performing electrophysiological recordings on recombinant GluN1-GluN2 receptor subtypes expressed in either *Xenopus laevis* oocytes or HEK293T cells. Interestingly, we found that YY-23 selectively reduced the glycine- and glutamate-induced maximal responses of GluN1-GluN2C and GluN1-GluN2D NMDARs in a dose-dependent manner (see IC_50_ values in Table S3). In contrast, YY-23 had minimal effects on both GluN1-GluN2A and GluN1-GluN2B receptors (Fig. 1b, c, f, g).

Next, we examined the effect of YY-23 on the current–voltage (I–V) relationship of GluN1-GluN2D receptors. Unlike conventional voltage-dependent magnesium blocks, the I–V curve exhibited a voltage-independent profile upon the application of 10 μM YY-23 (Fig. 1h). There was no significant difference in the inhibition level when the holding potential was set to −60, −40 or +40 mV (Fig. 1i). Next, to elucidate whether YY-23-mediated inhibition relies on the gating states of GluN1-GluN2D receptors, we applied 10 μM YY-23 to the nonactivated receptors with an equivalent duration as that used for the activated receptors. The results demonstrated that YY-23 also acted on the apo state of GluN1-GluN2D receptors and displayed a remarkable inhibitory effect (Fig. 1j). We subsequently examined YY-23-mediated inhibition at various doses of glycine or glutamate and found that the inhibition occurred in an agonist concentration-independent manner (Fig. 1k). Taken together, these data imply that YY-23 is an allosteric inhibitor rather than a competitive antagonist or a channel pore blocker of GluN1-GluN2D NMDARs.

### Structural determination of YY-23-mediated inhibition of GluN1-GluN2D receptors

Next, we tried to identify structural elements in NMDARs that might be responsible for the action of YY-23. GluN1 and GluN2 subunits share similar topological architectures, with two tandem extracellular domains comprising an N-terminal domain (NTD) and a ligand binding domain (LBD), a transmembrane domain (TMD) where the ion channel is located and an intracellular C-terminal domain (CTD) (Fig. 2a, b) (12). Both the NTD and LBD have a bilobed clamshell-like architecture, and the LBD is composed of S1 and S2 segments (Fig. 2a). We constructed truncated or chimeric subunits to search for the structural elements involved in YY-23-mediated inhibition (Fig. 2c, d).

**Figure 2.**
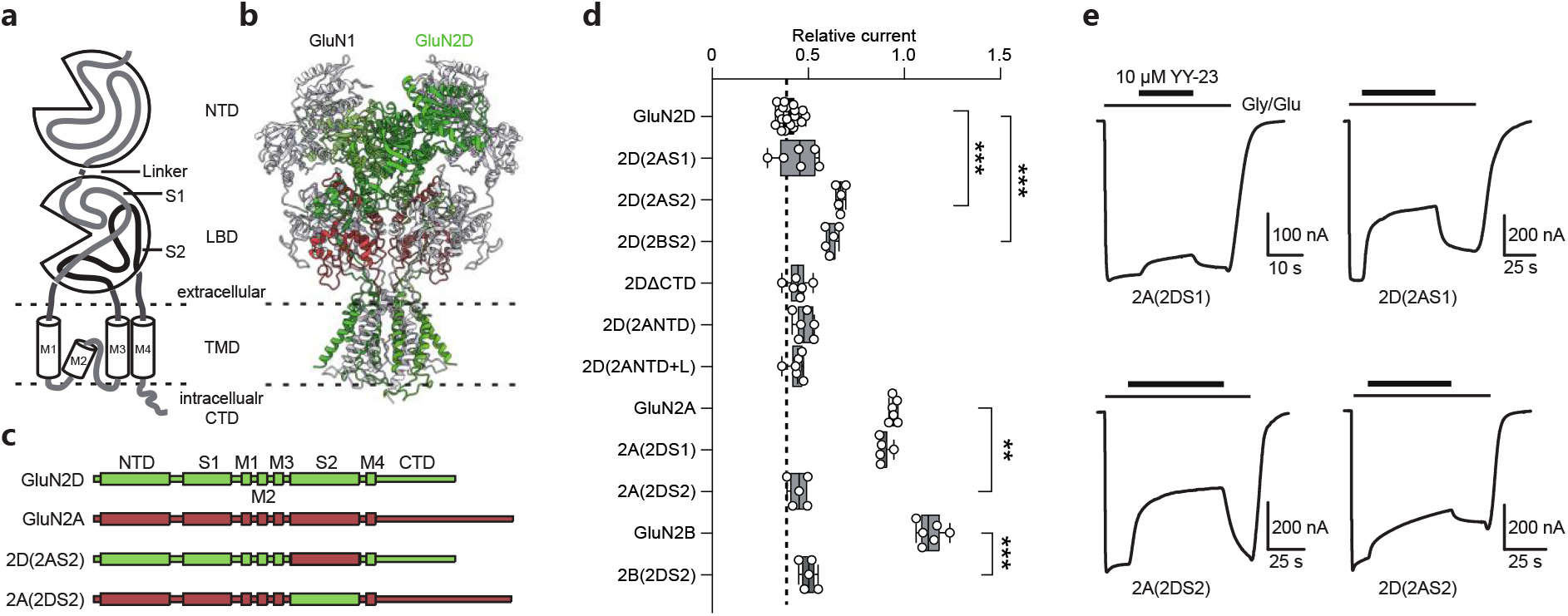
Structural determinant for subunit-selective YY-23 inhibition. (a) Cartoon illustration of the topological structure of GluN subunits with an N-terminal domain (NTD), a ligand binding domain (LBD) composed of S1 and S2 portions, a transmembrane domain (TMD) and an intracellular C-terminal domain (CTD). (b) Homologous model of the GluN1-GluN2D receptor structure built based on the GluN1-GluN2A cryo-EM structure (PDB code: 6ira) using SWISS-MODEL, with the S2 portion highlighted in brown. (c) Linear representation of GluN2A (brown), GluN2D (green) and chimeric subunits replacing the S2 portions between GluN2A and GluN2D subunits. (d) Responses to 10 μM YY-23 on wild-type receptors and all chimeric or truncated receptors expressed in Xenopus oocytes. The dashed line aligned with the median value of response of the GluN1-GluN2D wild-type receptor provides a benchmark for the YY-23-mediated inhibition effect. Note that replacing the S2 portion of GluN2A or GluN2B with the GluN2D S2 portion significantly increased YY-23 inhibition. The data were analysed by Brown-Forsythe ANOVA test followed by post hoc Dunnett’s T3 multiple comparisons. (e) Representative traces of chimeric receptor responses to 10 μM YY-23. S2 portion replacement had a gain-of-function effect on GluN1-GluN2A (2DS2) receptors and a loss-of-function effect on GluN1-GluN2D (2AS2) receptors; however, S1 portion replacement did not have these effects. In the box plot graphs, the bars show the min/max values, the boxes show the lower/upper quartiles, and the lines represent the median values. **p < 0.01 and ***p < 0.001.

Recording results showed that the modified GluN1-GluN2D receptors with the replacement of either the GluN2D-NTD, the NTD plus linker, or the S1 portion with that of the GluN2A subunit displayed no significant change in YY-23 inhibition level compared with that of GluN1-GluN2D wild type receptors (Fig. 2d). Notably, the GluN1-GluN2D (2A-S2) and GluN1-GluN2D (2B-S2) receptors showed intermediate declines in the inhibitory effect of YY-23 (Fig. 2d, e). More strikingly, GluN1-GluN2A (2D-S2) or GluN1-GluN2B (2D-S2) receptors displayed significant gain-of-function of YY-23-mediated inhibition (Fig. 2c, e). In contrast, the chimeric GluN2A subunit that had the S1 portion replaced with GluN2D showed no gain of YY-23-mediated inhibition (Fig. 2d, e). We further substituted individual residues in the TMD of GluN2D with their homologous amino acids in the GluN2A subunit or deleted the CTD of the GluN2D subunit. None of the mutant or truncated constructs led to a significant loss-of-function effect of YY-23 (Fig. S2a, b and Fig. 2d). Altogether, these data reveal that the S2 region of the GluN2D subunit is critical for the allosteric inhibition induced by YY-23.

### YY-23 inhibits native GluN2D-containing NMDARs in the mPFC

The GluN2C subunit is dominantly expressed in the cerebellum, so we selected GluN2D for the following *in vivo* studies, as it is widely distributed in the brain (20) and closely related to depression and anxiety (21, 22). However, very little is known about the cell type-specific expression and functional role of the GluN2D subunit in the adult cortex. We performed multiplex fluorescence in situ hybridization (FISH) to examine the mRNA expression of *Grin2d* at interneurons and pyramidal neurons in adult mice at 8 weeks (Fig. 3a-d). We found that *Grin2d* preferentially colocalized with cells positive for *slc32a1* (encoding GAD67) rather than those positive for *slc17a7* (encoding Vglut1). In addition, we analyzed mRNA expression level of five major subunits of NMDARs using single cell RNA-seq database. The result demonstrated the transcriptomic taxonomy tree of cell-type clusters organized in the cortex and hippocampus. In contrast to *Grin2a* and *Grin2b, Grin2d* is predominately distributed in the inhibitory neurons (Fig. S3). The results confirmed that the *Grin2d* subunit was expressed in the mPFC in adult mice and preferentially colocalized with interneurons rather than pyramidal neurons.

**Figure 3.**
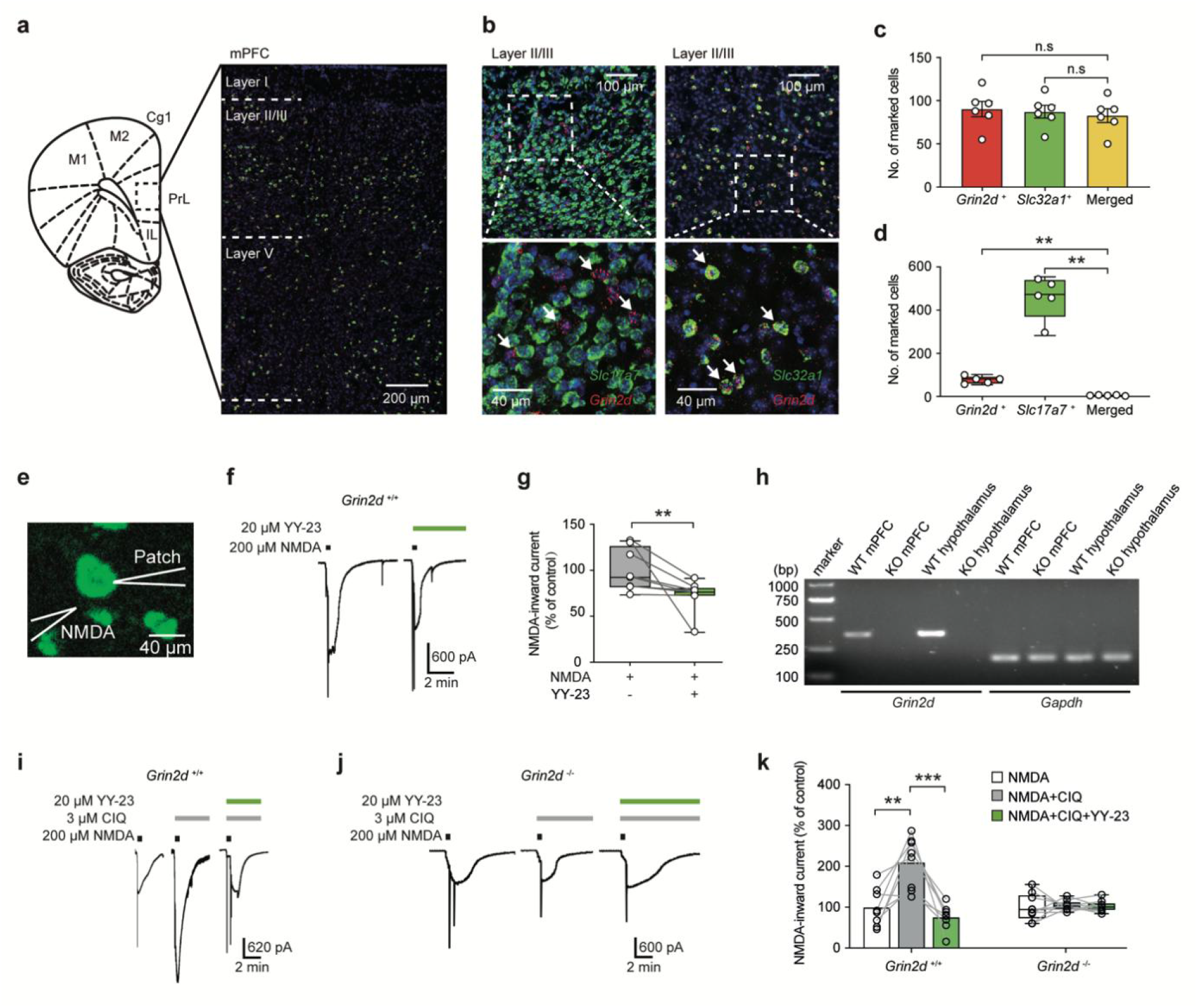
YY-23 abolished the CIQ potential of interneurons located in layer II/III of the mPFC. (a) Illustration of the coronal plane of the mouse brain sectioned in the left panel, with the mPFC region highlighted by a dashed box. The brain slice beside showing the confocal results, with DNA labelled with DAPI (blue) and the mRNA of Grin2d (red) and Slc32a1 (green) labelled with their probes through the multiplex fluorescence in situ hybridization (FISH) technique. Dashed lines distinguish the three layers of the mPFC in mouse brain. PrL, prelimbic cortex; IL, infralimbic cortex; Cg1, cerebral cortex, area 1. (b) Presentation of the mRNA localization of Slc17a7 (green, represents pyramidal neurons, left column), Slc32a1 (green, represents interneurons, right column) and Grin2d (red). The bottom panels in each column are the magnified images of the dashed boxes above. Cells with positive (+) single or double mRNA signals were counted (c and d). Note that 94.8% interneurons expressed Grin2d (highlighted) by white arrows, but only 1.2% pyramidal neurons expressed Grin2d. One-way ANOVA was used to analyse data with a normal distribution (c). Data with heterogeneity of variance (d) were tested by Brown-Forsythe ANOVA followed by post hoc Dunnett’s T3 multiple comparisons. (e) Illustration of patch-clamp recordings in a GAD67-GFP-positive brain slice containing the mPFC after agonist or drug administration. (f) Slice recording for the Grin2d^+/+^ mouse mPFC. YY-23 at 20 μM resulted in a significant decrease in peak amplitude, as shown in (g). (g) Box plots showing the peak amplitude induced by NMDA in the presence or absence of YY-23. Paired comparisons with abnormal data distributions were performed by the Wilcoxon test. (h) Reverse transcription-PCR (RT-PCR) of Grin2d gene transcription in mPFC and hypothalamus. WT represents wild type genotype and KO represents Grin2d-knockout. (i and j) mPFC slice recordings in Grin2d^+/+^ and Grin2d^-/-^ mice. Note that 20 μM YY-23 abolished 20 μM CIQ-induced potentiation in Grin2d^+/+^ mouse slices but not Grin2d^-/-^ mouse slices. (k) Box plots showing that the potentiation of peak amplitude induced by CIQ in Grin2d^+/+^ was blocked in Grin2d^-/-^mice. The peak amplitude in the absence of YY-23 and CIQ was set to 100%. Data with normal distribution and equal variability of differences (k in Grin2d^+/+^ slices) were tested by repeated–measurement one-way ANOVA followed by Tukey’s multiple comparisons. Data with heterogeneity of variance (k in Grin2d^-/-^ slices) were test by Geisser-Greenhouse correction. Each point represents a single cell. The bar graphs show the mean ± SEM. In the box plot graphs, the bars show the min/max values, the boxes show the lower/upper quartiles, and the lines represent the median values. *p < 0.05 and **p < 0.01.

Next, we hypothesized that YY-23 may preferentially act on GluN2D-containing NMDARs and then investigated the effects of YY-23 on the NMDAR activity of layer II/III interneurons in adult mPFC slices. NMDA-evoked inward currents of GAD67-EGFP-labelled interneurons were recorded by patch-clamp recordings (Fig. 3e). We found that YY-23 significantly inhibited NMDA-evoked currents (Fig. 3f, g). To further investigate whether YY-23 selectively acted on native GluN2D-incorporated NMDARs, we applied the selective allosteric potentiator CIQ to boost the currents of GluN2D-containing NMDAR currents (23). We found that YY-23 significantly blocked the enhancement of NMDA-evoked currents potentiated by 3 μM CIQ (Fig. 3i, k). However, in *Grin2d*-knockout (KO) mice (authenticated by quantitative PCR, Fig. 3h), neither CIQ nor YY-23 had any effect on NMDA-evoked currents (Fig. 3j, k). These results indicate that YY-23 selectively blocks the GluN2D subunit-mediated inward current in mPFC interneurons.

### The rapid antidepressant action of YY-23 is dependent on the GluN2D subunit

To assess whether YY-23 produced antidepressant-like effects, we first employed naïve mice to optimize the dose regimen for YY-23. YY-23 was acutely administered to naïve mice 30 min prior to the behavioural tests. In the novelty-suppressed feeding test and forced swimming test, YY-23 significantly decreased the feeding latency and immobility time in a dose-dependent manner (Fig. 4a, b). In addition, the total distance travelled did not show obvious differences in the open field test (Fig. 4c), indicating that no abnormal locomotor activity was induced by YY-23. These results showed that YY-23 produced obvious antidepressant-like effects upon acute administration.

**Figure 4.**
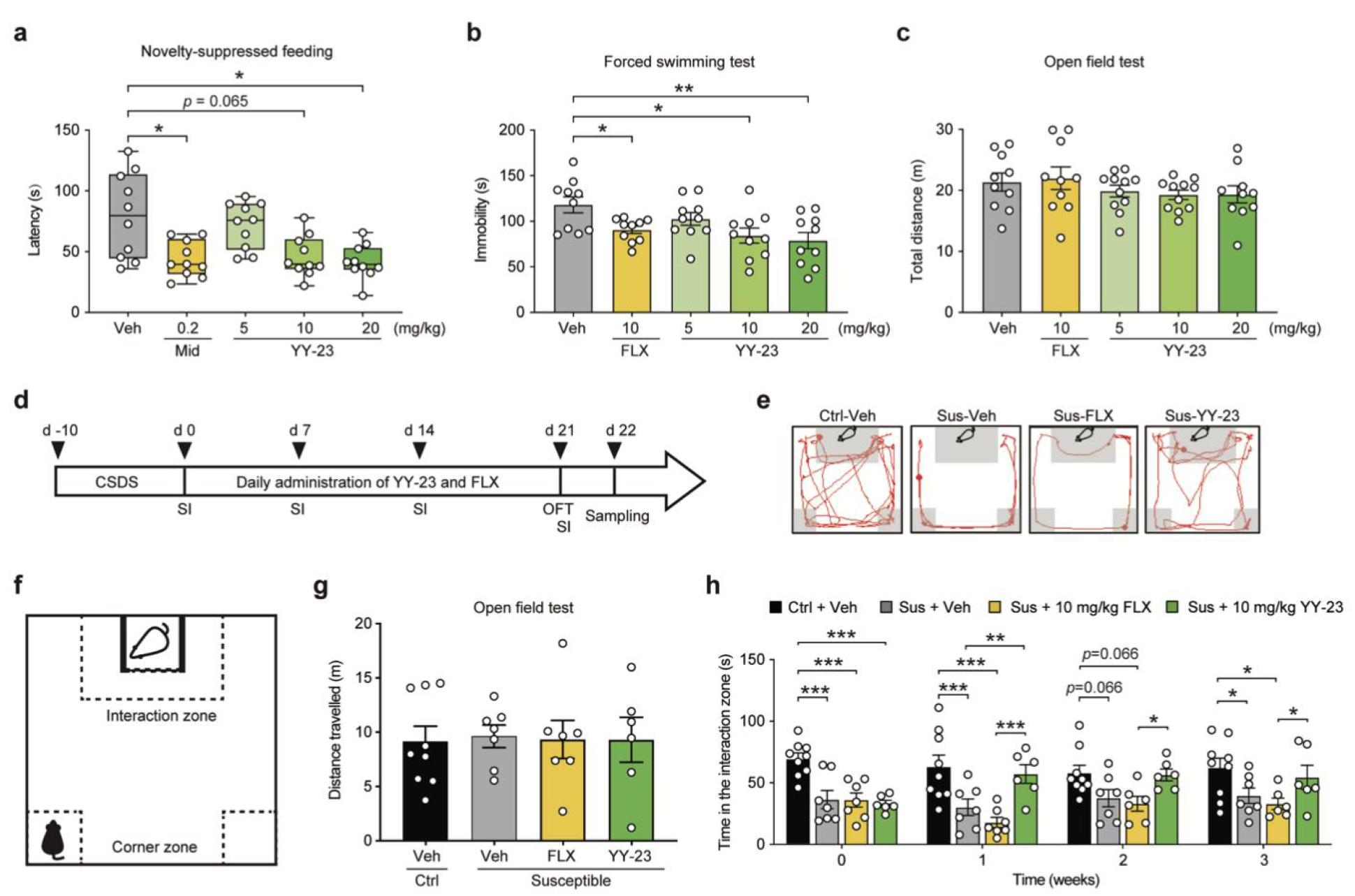
YY-23 produced a rapid antidepressant effect. YY-23 had dose-dependent antidepressant-like effects in the novelty-suppressed feeding test (a) and forced swimming test (b) but did not affect locomotion in the open field test (c). Data with heterogeneity of variance (a) were analysed by Brown-Forsythe test followed by post hoc Dunnett’s T3 multiple comparisons. Data with a normal distribution and equal variability (b and c) were tested by one-way ANOVA followed by Dunn’s correction for multiple comparisons. (d) Timeline for behavioural tests, drug administration and brain tissue sampling in the depressive model of chronic social defeated stress (CSDS). After 10 days of social stress exposure, mice that exhibited social phobia were chronically treated with vehicle, fluoxetine (FLX) or YY-23. Naïve animals were treated with vehicle as a control group. Social interaction (SI) tests were performed once a week. Twenty-four hours after the last drug administration, the tissues were sampled. (e) The red lines represent the exploratory traces of C57BL/6N mice travelling in the arenas after 21 days of drug administration. Note that YY-23 significantly improved the exploration of the interaction zone (grey box), but fluoxetine did not. (f) Illustration of the social interaction test paradigm in the presence of the target CD-1 mouse. The defeated mouse (C57BL/6N) was initially set at the corner zone (dashed box at the bottom corner) to begin its free exploration. The dashed square at the top was defined as the social interaction zone. (g) On Day 0 after CSDS, there were no significant differences among groups in the open field test, excluding disturbance of locomotor activity. The data were tested by one-way ANOVA. (h) Statistical analysis of the time spent in the interaction zone. Note that social interaction in the animals administered YY-23 but not fluoxetine was rescued within one week. Sus represents susceptible mice. The data were analysed by a mixed-effects model with the two-stage linear step-up procedure of Benjamini, Krieger and Yekutieli for multiple. The bar graphs show the mean ± SEM. In the box plot graphs, the bars show the min/max values, the boxes show the lower/upper quartiles, and the lines represent the median values. *p < 0.05, **p < 0.01 and ***p< 0.001.

We next developed a mouse depressive model of chronic social defeat stress (CSDS) (24) to determine whether YY-23 prevented depressive phenotypes induced by stress (Fig. 4d-f). A social interaction (SI) ratio equal to 1, in which equal time is spent in the presence versus absence of a social target (CD-1), was used as the threshold for dividing defeated mice into the susceptible and resilient categories. Mice below this criterion were grouped as susceptible. After 10 days of the CSDS procedure, approximately 60% of stressed mice showed defeated phenotypes with the reduced time spent in the interaction zone (Fig. 4e, h) and showed normal locomotor activity compared with control mice tested by the open field test (Fig. 4g). All susceptible mice were then randomly divided into three groups and treated with vehicle, 10 mg/kg YY-23 or 10 mg/kg fluoxetine. The control group was administered vehicle. Depressive-like behaviour was examined according to the time spent in the interaction zone when aggressive target mice were present. After one week of daily YY-23 administration, stressed mice and control mice spent similar amounts of time in the interaction zone, indicating that the depressive-like symptoms had recovered to normal levels. In contrast, 3 weeks of fluoxetine treatment did not reverse social defeat behaviours (Fig. 4h). This finding was also similar with our previous results (16), which were examined in the depressive model of chronic unpredicted mild stresses (CUMS). The results suggest that YY-23 reverses depressive phenotypes induced by CSDS and has a faster antidepressant onset than fluoxetine.

Next, by using *Grin2d* knock out (*Grin2d^-/-^*) and heterozygous (*Grin2d^+/-^*) mice, we examined whether inhibition of GluN2D-containing NMDARs by YY-23 is the molecular trigger for the antidepressant effects. In the forced swimming and tail suspension tests (Fig. 5a, b), we found a significantly decreased immobility time after a single dose of YY-23 than after treatment vehicle in the wild type mice; however, these effects of YY-23 were abolished in the *Grin2d* knock out and heterozygous mice. The locomotor activity, including total distance travelled (Fig. 5c) and central distance travelled (Fig. 5d), were unchanged in the *Grin2d^+/+^, Grin2d^+/-^* and *Grin2d^-/-^* mice. These findings suggest that GluN2D-containing NMDARs are the targets for the antidepressant effects induced by YY-23.

**Figure 5.**
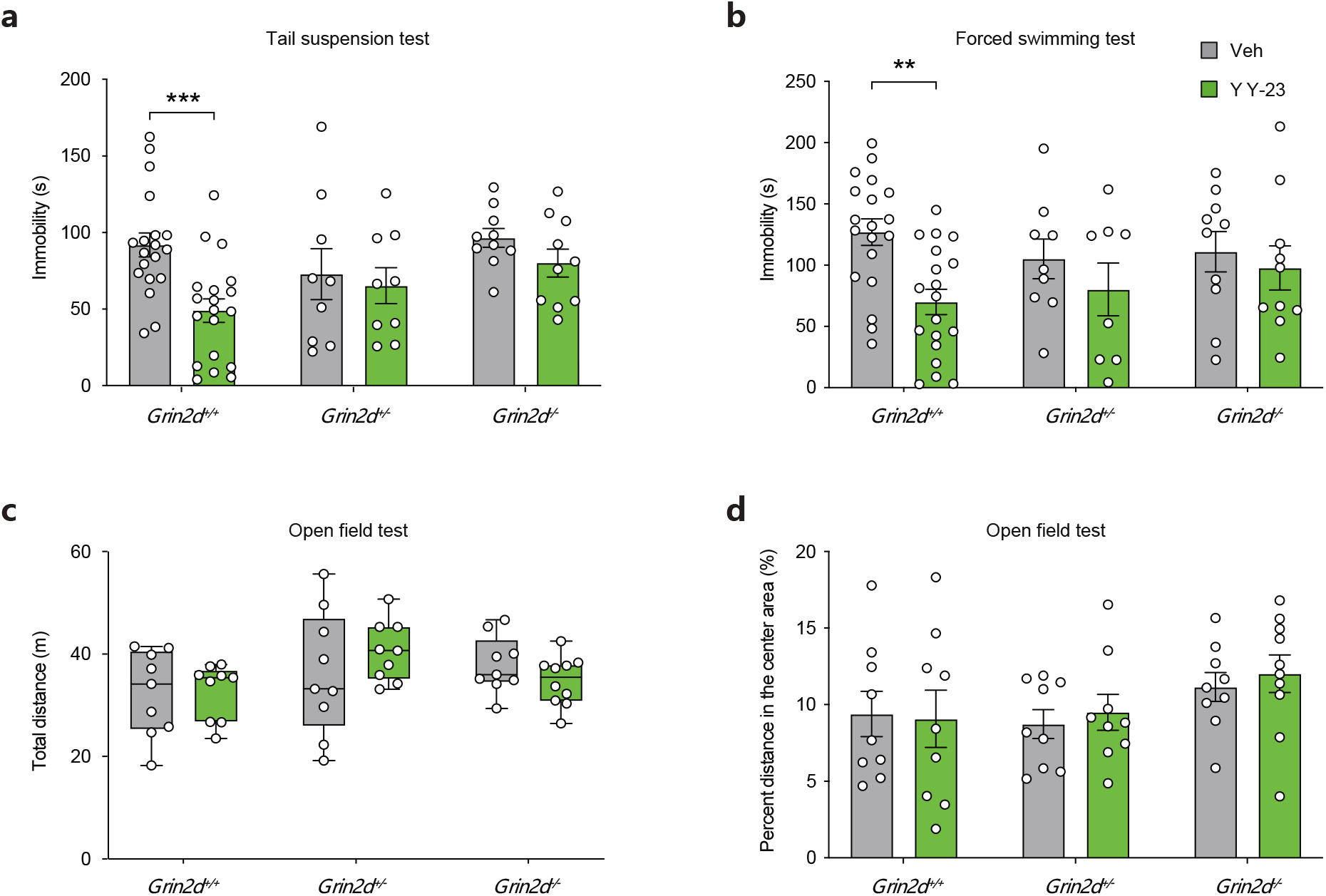
Genetic deletion of Grin2d abolished the antidepressant-like effect of YY-23 in animal models. In the Grin2d^+/+^ mice, YY-23 significantly decreased the immobility time in both the forced swimming test (a) and tail suspension test (b), which were prevented in the Grin2d-knockout (Grin2d^-/-^) and heterozygous (Grin2d^+/-^) mice. Both YY-23 administration and Grin2d knock out (Grin2d^+/-^ and Grin2d^-/-^) did not affect the total distance travelled in the open field test (c and d). Data with a normal distribution were tested (a, b and d) by two-way ANOVA followed by Bonferroni correction for multiple comparisons. Data with an abnormal distribution were tested (c) by Friedman’s two-way ANOVA. The bar graphs show the mean ± SEM. In the box plot graphs, the bars show the min/max values, the boxes show the lower/upper quartiles, and the lines represent the median values. **p < 0.01 and ***p< 0.001.

### Inhibition of GluN2D-containing NMDARs on GABAergic interneurons enhances excitatory transmission

To test whether YY-23 exerted functional effects on mPFC interneurons, we first recorded action potentials (APs) via a series of current injections in mPFC GAD-67-EGFP-positive interneurons (Fig. 6a-c). The numbers of APs were significantly reduced by YY-23 at concentrations of 1, 10 and 30 μM in a concentration-dependent manner (Fig. 6b, d, e). Given that YY-23 weakened the excitability and firing output of interneurons, we wonder how these changes might affect inhibitory output onto nearby pyramidal neurons. Thus, we compared spontaneous inhibitory postsynaptic currents (sIPSCs) in the mPFC layer V pyramidal neurons in WT and KO mice. YY-23 at 10 μM resulted in a decreased in frequency but not amplitude of sIPSC in WT mice (Fig. 6f-h and l-m). This observation also occurred in the miniature inhibitory postsynaptic currents (mIPSC) recorded in the pyramidal neurons (Fig.S4 a-h). The results indicate that the GABAergic output of interneurons into pyramidal neurons was inhibited. As sIPSC makes postsynaptic more likely to inhibit firing of action potential, we then examined whether this inhibition affected the excitability of pyramidal neurons. The results demonstrated that YY-23 (10 μM) increased the number of action potentials (Fig. 6n-q) and miniature EPSC (mEPSC) frequency and amplitude (Fig. S4 i-q), suggesting that YY-23 may enhance the excitability of pyramidal neurons by disinhibiting GABAergic output of interneurons. Furthermore, we found that YY-23 induced-inhibition of sIPSC and enhancement of action potential in the pyramidal neurons were prevented in the *Grin2d* mice (Fig. 6i-k and l-m), suggesting that GluN2D subunit may contribute to YY-23-induced reduction of inhibitory output onto nearby pyramidal neurons and enhancement of excitability of pyramidal neurons.

**Figure 6.**
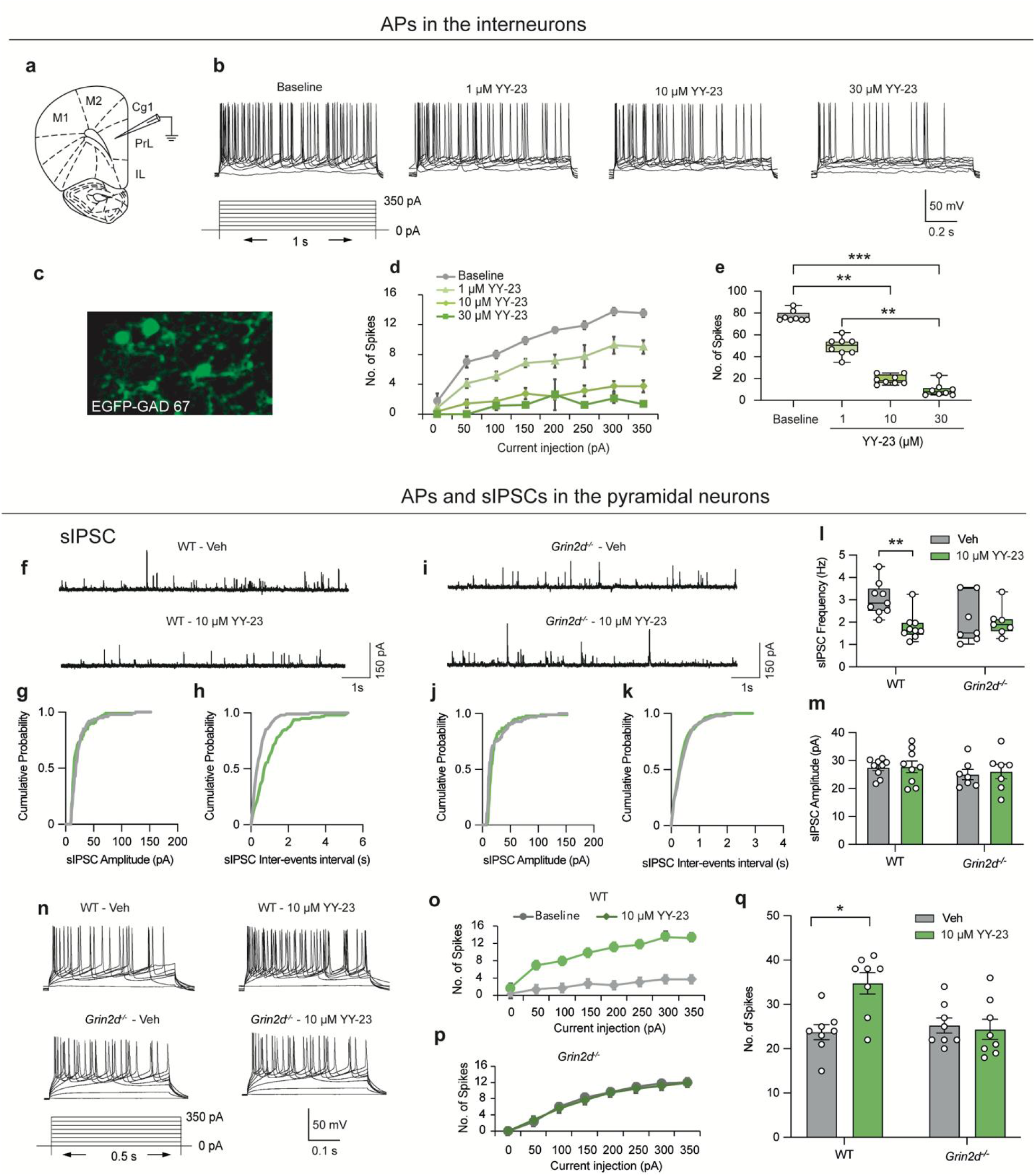
YY-23 upregulated pyramidal neuron activity by inhibiting interneuron activity. (a-e) Action potentials recorded in the mPFC interneurons. Schematic diagram of the coronal plane of a mouse brain sectioned at bregma 1.54~2.10 mm. The recoding electrode was placed at the prelimbic area (Prl) of the mPFC (a). Illustration of GAD-67-EGFP-positive interneurons in the mPFC (c). Representative active potentials (b) and response curves between current injections and the number of spikes (d) evoked by step current injections ranging from 0~350 pA in 50 pA increments in the presence and absence of YY-23. Box plot graphs showing that YY-23 significantly inhibited APs in a concentration-dependent manner (e). The data with abnormal distribution were analysed by Kruskal–Wallis one-way ANOVA followed by Dunn’s correction for multiple tests. (f-m) sIPSCs recorded in the mPFC pyramidal neurons. Representative sEPSC traces in the wt (f) and Grin2d ko (i) mice after vehicle and 10 μM YY-23 treatment. Cumulative distribution of sEPSC amplitudes (g, j) and interevent intervals (h, k). The data were analysed by Kolmogorov–Smirnov test. The box plot graph (l) show frequency and bar graphs (m) show average amplitude after the application of vehicle and YY-23 in the wt and ko mice. Data with abnormal distribution were analysed by wilcoxon matched-pairs signed rank test (l). Data with normal distribution were analysed by multiple paired t test (m). (n-q) Action potentials recorded in the mPFC pyramidal neurons. Representative APs traces (n) and response curves between current injections and the number of spikes (o, p) of mPFC pyramidal neurons evoked by step current injections ranging from 0~350 pA in 50 pA increments in the presence and absence of YY-23. (q) Bar graph shows average APs after the application of vehicle and YY-23 in the wt and ko mice. Data with abnormal distribution were analysed by multiple paired t test. The bar graphs show the mean ± SEM. The box plot graphs show the min/max values, the boxes show the lower/upper quartiles, and the lines represent the median values; the data were analysed by Wilcoxon tests. *p < 0.05, **p < 0.01 and ***p < 0.01.

Given that the SSRI fluoxetine altered neither the mEPSC frequency (Fig. S5 a-f) nor the amplitude (Fig. S5 a-c and g-i) in pyramidal cells, suggested a unique neuropharmacological mechanism of YY-23 compared with SSRIs. The results implied that YY-23 directly inhibits GluN2D-containing NMDARs within interneurons, which reduces the inhibitory output of GABAergic interneurons and in turn results in the disinhibition of pyramidal cells. Thus, we speculate that the fast antidepressant effect of YY-23 is likely related to the induced enhancement of excitatory synaptic transmission in the mPFC in a characteristic manner involving disinhibition.

### Proteomics analysis of the mPFC network and synaptic plasticity in response to YY-23

To further investigate the pathophysiological mechanism underlying stress-induced depression and to identify the phenotype-specific molecular function of YY-23, we employed quantitative proteomics to analyse protein expression in the mPFC. Three animals were used per group for brain tissue sampling 24 h after 3 weeks of drug administration (Fig. 4d), and a total of 4915 distinct proteins were quantified by mass spectrometry. Proteomic analysis revealed that the expression levels of 120 proteins were significantly different between the susceptible and control groups. Unsupervised hierarchical clustering analysis of these 120 proteins clustered them into four distinct groups representing the four different responses to the treatments (Fig. 7a). We also found that the changes in 26 altered proteins in the susceptible group (stress *vs*. ctrl) were significantly reversed by YY-23 (YY-23 *vs*. stress, Fig. 7b). Thus, these proteins represent some common components that are oppositely regulated by stress and YY-23. The clustering analysis results shown in the heatmap further revealed that the 26 proteins, including 11 proteins that were–upregulated and 15 that were downregulated by stress, could be regulated in an opposite manner by YY-23 (Fig. 7c). In contrast, fluoxetine, which is an antidepressant with a long response lag, rescued the levels of only 12 of 120 proteins that were changed by stress (Fig. S6).

**Figure 7.**
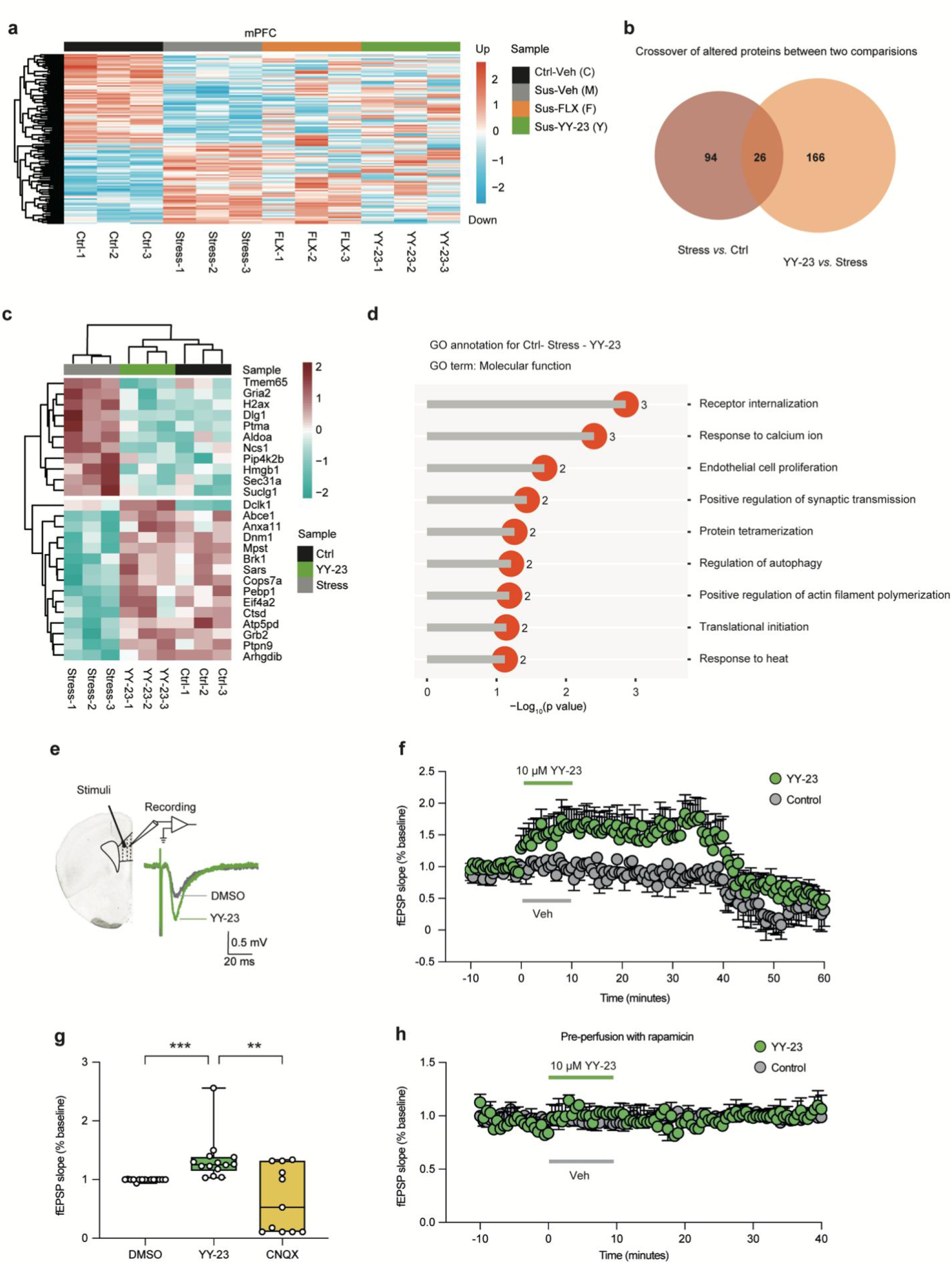
YY-23 enhanced synaptic transmission according to proteomic analysis and the LTP test. (a) Clustering analysis of differentially expressed proteins in the mPFC from the control group (Ctrl-Veh) and three groups of susceptible mice administered vehicle, 10 mg/kg fluoxetine (Sus-FLX) or 10 mg/kg fluoxetine (Sus-YY-23) (n=3 for each group). The sampling strategy is shown in Fig. 4d. (b) Venn diagram revealing the number of altered proteins in the stress group vs. ctrl group and in the YY-23 group vs. stress group. Note that the administration of YY-23 in susceptible mice rescued the levels of 26 proteins. (c) Clustering analysis of 26 proteins showing that the levels of the altered proteins in the susceptible mice (Stress 1-3) were rescued by the administration of YY-23. (d) GO enrichment analysis showed the biological processes associated with the proteins shown in (c). (e) Schematic diagram of fEPSP recording and example traces. The stimulating electrode on the left was placed on layer VI of the PFC slice, and the recording electrode was placed on layer II/III. (f) fEPSP slopes were recorded with DMSO as a control or with 10 μM YY-23 application (Vehicle or YY-23 was applied at 0 minutes and lasted 10 mins). CNQX was applied at 40 min to detect the participation of AMPA receptors. The data were analysed by Kolmogorov–Smirnov test. (g) Box plot graph show that YY-23 significantly increased the fEPSP slope but that this effect was abolished by CNQX. Data with abnoumal distribution were analysed by Kruskal–Wallis one-way ANOVA. (h) fEPSP slopes were recorded with DMSO as a control or with 10 μM YY-23 application pre-perfusion with rapamycin. **p < 0.01and ***p< 0.001.

To gain insight into the involvement of these proteins in different molecular and biological functions, GO annotation of the 26 proteins inversely regulated by YY-23 was performed. Nine terms were significantly enriched in the molecular function (MF) category. The identified proteins were mainly involved in receptor internalization, response to calcium ions and positive regulation of synaptic transmission (Fig. 7d). The results of proteomics analyses and the findings that YY-23 enhanced excitatory neurotransmission in the mPFC (Fig. 6) suggest a close association between the identified proteins and synaptic transmission after treatment with YY-23.

To further detect the underlying synaptic modulation by YY-23, we recorded field excitatory postsynaptic potentials (fEPSPs) in the mPFC. We placed the stimulating electrode on layer V and the recording electrode on layer II/III (Fig. 7e). Unexpectedly, the initial slope of the fEPSP was dramatically increased by the application of YY-23 within 10 min, and the effect lasted up to 40 min. The fEPSPs were inhibited by application of the α-amino-3-hydroxy-5-methyl-4-isoxazolepropionic acid (AMPA) receptor antagonist CNQX, confirming that the fEPSPs were AMPA receptor-dependent (Fig. 7f, g).

We wonder what the mechanism is underlying YY-23-induced long-last enhancement in excitatory synaptic transmission. Studies demonstrate that mammalian target of rapamycin (mTOR) signalling pathway localized to dendrites and cell body and regulated translation, have been linked to fast antidepressant and long-term synaptogenesis(25, 26). Here, we pre-treated the mPFC slice with selective mTOR inhibitor rapamycin at 5μM for 30 min and then examined the fEPSP again. The result showed that the enhancement of fEPSP was prevented by pretreatment of rapamycin at 5 μM (Fig. 7h). These findings suggest that induction of long-lasting enhancement in excitatory synaptic transmission and antidepressant activities may be dependent of mTOR signaling pathway.

## Discussion

In the present study, we identified a timosaponin derivative, YY-23, that selectively inhibited the activity of GluN2C- and GluN2D-containing NMDARs. Electrophysiological recordings further suggested that YY-23 neither directly blocked the channel gate nor competed with agonist binding, presumably acting as a negative allosteric modulator of GluN1-GluN2D receptors. More importantly, YY-23 produced fast antidepressant effects in mouse models that were abolished in *Grin2d*-KO mice. In the mPFC region of the mouse brain, GluN2D was predominantly expressed at GABAergic interneurons. YY-23 directly inhibited native GluN2D-containing NMDAR currents in the mPFC and decreased the output of GABAergic interneurons, subsequently disinhibiting the excitability of pyramidal cells and enhancing synaptic plasticity.

GluN1-GluN2D receptors display distinct biophysical properties, with the highest agonist potency, the longest exponential deactivation time course, and the lowest channel open probability among NMDAR subtypes (12, 14). The activity of GluN1-GluN2D receptors can be allosterically regulated by endogenous protons (27); neurosteroids (28); and exogenous allosteric compounds, including CIQ 23, QNZ46 (29) and DQP-1105 (30). Allosteric modulators, which bind to sites other than orthosteric agonist-binding sites (31), usually show subunit-specific actions without interfering with basic channel function (32, 33). For instance, allosteric modulators such as rapastinel generally display fewer side effects than the channel pore blocker ketamine, which usually exhibits psychotomimetic or dissociative effects (34). Here, we demonstrated that the mechanism of action of YY-23 depended on the involvement of the GluN2D-S2 segment in the LBD, which has been previously proposed to have a significant influence on the actions of QNZ46 (18), DQP-1105 (30) and neurosteroids (35). These results indicate that the S2 region is a promising target for the development of subtype-selective allosteric modulators.

Preclinical and clinical studies have characterized the neuroanatomical and functional abnormalities of the PFC in MDD models and patients (36–40). The PFC is vulnerable to stress, and even short and mild stressors can cause rapid dendritic shrinkage and dramatic prefrontal cognitive disabilities (41, 42). Interestingly, aberrant gene expression and dysfunction of NMDARs have been found in the PFC in MDD patients (43–45). A large number of studies have demonstrated that chronic stress gives rise to excitation/inhibition (E/I) imbalance in the PFC (46–50), which is strongly associated with depression. This has been further evidenced by the discovery that drugs targeting NMDARs can rectify the E–I imbalance. For example, ketamine has been proposed to act on NMDARs at GABAergic neurons (51), while the PAM rapastinel targets NMDARs at glutamatergic neurons (15). Although both drugs trigger different cell type-specific regulation of NMDARs, they ultimately exhibit promising antidepressant actions (15).

Consistent with the E–I balance theory, we found here that YY-23 precisely targeted GluN2D-containing subtype NMDARs at GABAergic interneurons. It reduced the inhibitory input (decreased APs and IPSC) to pyramidal cells and indirectly enhanced excitatory transmission in the mPFC (Fig. 6, 7 and Fig. S3). Based on previous studies, this pyramidal cell disinhibition triggered by NMDAR antagonists might activate downstream signalling, including phospho-mTOR, BDNF, and excitatory transmission mediated by glutamatergic AMPA receptors (AMPARs), which will help reestablish synaptic communication that is impacted in depression (52, 53). Here, we confirmed that YY-23 induced upregulation of excitatory synaptic transmission (Fig. S3 i-q) and long-term potentiation (AMPAR-mediated fEPSPs, Fig. 7f) in the mPFC. Moreover, the long-term potentiation and antidepressant actions of YY-23 could be blocked by the rapamycin, suggesting mTOR signaling may be involved in the mechanism of antidepressant effects of YY-23. Together, these results demonstrated that GluN2D triggered pyramidal cell disinhibition and that reestablishment of synaptic communication underlies the antidepressant actions of YY-23. More importantly, our study provides insights for the development of therapeutic antidepressant agents targeting GluN2D-containing NMDARs.

## Acknowledgments

Financial supports are gratefully acknowledged for the General National Natural Science Foundation of China (No. 31771115 and No. 31671049) to Shujia Zhu and Yang Li, respectively. The Key Program of National Natural Science Foundation of China (No. 81030065) to Chenggang Huan. The National Key R&D Program of China (2017YFA0505700), the Strategic Priority Research Program of the Chinese Academy of Sciences (XDB32020000), the Shanghai Municipal Science and Technology Major Project (2018SHZDZX05) to Shujia Zhu. The Strategic Leading Science and Technology Project of the Chinese Academy of Sciences (XDA12040214), the “Personalized Medicines—Molecular Signature-based Drug Discovery and Development”, Strategic Priority Research Program of the Chinese Academy of Sciences (XDA12040220) and the National Major Scientific and Technological Special Project for “Significant New Drugs Development” (201909301102) to Yang Li. The Science and Technology Commission of Shanghai Municipality (No. 184319071000 and No. 19140903102) to Yang Li and Fei Guo, respectively. The Youngth Program of National Natural Science Foundation of China (No. 81803828) to Fang Liu. The Youth Innovation Promotion Association of Chinese Academy of Science (No. 2019280) Xiaoting Tian.

