## Supplementary material for "A novel antidepressant acting *via* allosteric inhibition of GluN2D-incorporated NMDA receptors at GABAergic interneurons": Supplemetnal information

### Supplementary Methods and Materials

#### *Animals*

Adult male C57BL/6N mice aged 8-12 weeks were used for electrophysiological recording and behavioural tests. Adult male CD-1 mice aged 4-6 months were used as aggressors to develop a depressive model of chronic social defeated stress (CSDS). The mice were raised under stable conditions (except for the development of stressed mice) with access to food and water *ad libitum*. All animal studies and experimental procedures were approved by the Animal Care Committees of Shanghai Institute of Materia Medica and the Center for Excellence in Brain Science and Intelligence Technology, Chinese Academy of Sciences. The experiments were carried out in accordance with EU Directive 2010/63/EU on the protection of animals used for scientific purposes. Double-blind experiments with drug administration were conducted, and the animals were randomly divided into groups.

#### *Transgenic mice construction and validation*

*Grin2d*-knockout mice were generated with a CRISPR–Cas9 system at Biocytogen Pharmaceuticals (Beijing) Co., Ltd. by deleting exon 2 of the *grin2d* gene from the mouse genome according to Ikda K *et al.* (1). In brief, a pair of sgRNAs (sgRNA 1: 5' AGCCTGCGCCGCTCCGCATG 3'; sgRNA 13: 5' TGGCAGAATGAGCCTAGTCT 3') was designed targeting the exon 2 segment of the *grin2d* gene. The deletion of exon 2 terminates the protein translation in exon 3 via frameshift mutation. To investigate the off-target effects that might be caused by the CRISPR/Cas9 system, we retrieved possible off-target sites for sgRNA1 and sgRNA13 using CCTop, a CRISPR–Cas9 target online predictor (https://cctop.cos.uni-heidelberg.de:8043), with the default parameters and obtained 9 and 10 candidates for sgRNA1 and sgRNA13 (Fig. S7a), respectively. Next, we amplified DNA segments covering the possible gRNA recognition sites in the genomes of both wild-type and *grin2d*-knockout mice using the following primers (Fig. S7b):

| Candidates | Forward primer 5'-3' | Reverse primer 5'-3' |
|---|---|---|
| gRNA 1-1 | CTGTGATGGTAGGGGCACTG | CAGAAGAAGCACCCGAGGTT |
| gRNA 1-2 | ACCGAAATGCTAATACAAGCGG | GGTAACCGCCCAGTTGTTGT |
| gRNA 1-3 | GGTGTTTTCCGCTCTCCCAT | TCAGCTCATGTGCCAACTGT |
| gRNA 1-4 | CAATATGTGTCAGGCATCAGGT | TGTCCAAGACTGAGAACCTGT |
| gRNA 1-5 | CAGTTGACAGTGGTTTGCGT | GCAGACAACCAATGAGCACC |
| gRNA 1-6 | TGAGCTCTTCAACTGCTCCG | CCCAGTCACCAAGCCCTAAG |
| gRNA 1-7 | CGCTCCCCATGCGGATCTGCTC | GAGGCAGTTTGGAAGACGCA |
| gRNA 1-8 | TGCAGCCGCATTAAGGTGTTAT | TCAATAGCAGATGACAGCAG AGCA |
| gRNA 1-9 | TCCCTCCAGGTGCTATGGAA | TTTCCGTTTACGCTGTGGGT |
| gRNA 13-1 | ACAACTGTACCAGCCACAGG | GTGCAGCAAGCCATATCTGC |
| gRNA 13-2 | TGAGGACTCAGCTCTTCCCA | TCCAAGCCCTGAATGTCTGT |
| gRNA 13-3 | GCCCCGTACCTGTTTGATGGAAGAT | AAACCAGCAAAGGGCACCTA CTAGC |
| gRNA 13-4 | CACAGTACCTAAGTGCATAC | GAACTCCATGTACCTCTCTAG |
| gRNA 13-5 | ATACCCAAAGCGTGCAAAGG | AGAATCTGGCTGGGGATGGA |
| gRNA 13-6 | CAGGGGCTTTCTTTGTAGCC | TGCAGCCTTCTAGTTCACCA |
| gRNA 13-7 | CTGAGCTCTGCCTGTTCTTTG | GAGTGACCAACCAATGACTG |
| gRNA 13-8 | CAGCATCGGTGCAGCTGTTTC | AGCCATCTGGCTGAGTAATGG CTAA |
| gRNA 13-9 | GAGCCACATCTCAGGTCCCTG | GTCTCAGTGGGACACCAACTA TGCC |
| gRNA 13-10 | TCCAGCTACCAGTGTCTCAG | ATCATTCCCTTGCATGCCCA |

The PCR products were treated with End Prep Enzyme Mix for end repair, 5' phosphorylation and dA tailing in one reaction and then subjected to T-A ligation to add adaptors to both ends. Each sample was then amplified by PCR using the primers P5 (5' AGATCGGAAGAGCGTCGTGTAGGGAAAGAGTGT 3') and P7 (5' AGATCGGAAGAGCACACGTCTGAACTCCAGTCAC 3'). Both primers carried sequences that could anneal with the flow cell to perform bridge PCR and carried indexes to enable multiplexing. Then, equal amounts of PCR products were pooled and sequenced using an Illumina NovaSeq platform (Illumina, San Diego, CA, USA). The sequencing data were processed with CRISPResso2 for quantification of insertions and deletions at each possible off-target site. The editing efficiency was calculated as the number of reads containing mutations or indels divided by the total number of reads (Fig. S7c).

To further validate the gene expression of the *grin2d*-knockout mice, we detected the mRNA expression level of *grin2d* by quantitative PCR. We extracted total RNA (Kit No. B511311, Sangon Biotech) from the brain medial prefrontal cortex (mPFC) and hypothalamus tissues of wild-type and mutant mice and constructed a cDNA library using a cDNA reverse transcription kit (Kit No. 18080044, Invitrogen). Then, we amplified *grin2d* (forward primer: 5' GTGCTGGAGGAGTACGACTG 3', reverse primer: 5' GCAGAAGAAGTGGTTCCCCA 3', used for amplifying a segment of *grin2d* exon 2) and a housekeeping gene, *gapdh* (as a control, forward primer: 5' TGTGTCCGTCGTGGATCTGA 3', reverse primer: 5' TTGCTGTTGAAGTCGCAGGAG 3'), and analysed all the PCR products by agarose gel electrophoresis (Fig. 3h).

*HPLC–MS/MS characterization of the synthetic products*

Timosaponin BIII was isolated from *Rhizoma Anemarrhenae*, which was purchased from the medicinal materials market of Shanghai (Shanghai City, China). HPLC-grade acetonitrile was purchased from Dikma Company (Dikma, USA). The other reagents used were of analytical grade (Sinopharm Chemical Reagent Co. Ltd., China). Optical rotations were measured with a Perkin-Elmer 343 polarimeter. NMR spectra were measured on a Varian Mercury-400 (Varian, USA) using deuterium pyridine as the solvent and TMS as an internal standard. HR-ESI-MS was carried out on a Micromass TOF Ultima (Micromass, UK) in negative electrospray (ESI-) ionization mode. Semipreparative HPLC was performed on a Unimicro Technologies System (quaternary pump, China) with a Grace Apollo C18 (10 mm i.d. × 250 mm, ODS, 5 μm) semipreparative HPLC column including an EasyGuard Kit C18 (4 mm × 2 mm) guard column; the detector was a UV detector.

*Screening of pharmacological targets for YY-23 based on monoaminergic receptors*

We used a FLIPR® Calcium 4 Assay Kit (Molecule Devices) for the measurement of agonist- and antagonist-stimulated calcium signalling in cells (CHO-K1, Chempartner) expressing G-protein-coupled receptors (GPCRs), including $5HT_{2A}$, $5HT_{2B}$, and $5HT_{2C}$. The activity levels of these GPCRs can be functionally assessed using FLIPR-based fluorescence assays (2, 3). Here, we chose the fluorescence intensity to represent the activation and inhibition of GPCRs after agonist, antagonist or YY-23 application. We seeded 50 μL of cell suspension at the proper density into each well in 384-well assay plates (Corning, Cat. No. CC3712) and then incubated the cells in an incubator at 37 °C and 5% $CO_2$ for 16-24 hours. Next, we removed the culture medium from each of the plate wells and loaded 30 μl of calcium-sensitive dye (Component A, Molecular Devices) per well before incubating the plate for 1 hour in an incubator. The compounds to be tested were transferred to the compound plate with 30 μl of assay buffer (1× HBSS +20 mM HEPES, pH 7.4). The compounds were then added to the cell plate at 15 μl/well and incubated for 10 minutes at room temperature. Agonist was added at 22.5 μl/well, and the calcium flux signal was measured with FLIPR.

We used a Lance Ultra cAMP kit (Molecule Devices) to test the effects of compounds on $5HT_{1A}$,

$D_1R$, $D_2R$ and $D_4R$ in agonist and antagonist mode. The cells were cultured at 37 °C and 5% $CO_2$ with culture medium. Then, 100 nL of each compound was transferred to an echo machine. The cells were collected with stimulation buffer (1× HBSS with 5 mM HEPES+0.05 mM IBMX+0.1% BSA), plated at a proper density, centrifuged at 600 rpm for 3 min and then incubated with the compounds at room temperature for 60 min. After that, 5 μL of 4X Eu-cAMP tracer solution and 5 μL of 4X ULight™-anti-cAMP solution were added to the cells. The mixtures were centrifuged at 600 rpm for 3 min and incubated for 60 min. The cAMP signal was detected with Envision.

*Plasmid construction*

pcDNA3.1-based expression plasmids encoding wild-type rat GluN1-1a (GenBank accession number U08261, hereafter GluN1), GluN2A (GenBank accession number D13211), GluN2C (GenBank accession number M91563), GluN2D (GenBank accession number L31611) and mouse GluN2B (GenBank accession number BC172745) were generously provided by Dr. Pierre Paoletti (Institut de Biologie de l'Ecole Normale Supérieure, France). The amino acid numbering started from methionine in the signal peptide. We designed all the chimeric and truncated constructs based on segmenting of the GluN2 subunits. Briefly, the N-terminal domains (NTD) were segmented at Ala26-Trp390 for GluN2A and Gly39-Trp413 for GluN2D. The linkers (L) between the NTD and agonist binding domain (ABD) were segmented at Pro391-Asn404 for GluN2A and Ser414-Gln427 for GluN2D. ABD S1 was segmented at His405-Ser556 for GluN2A, His405-Asp557 for GluN2B and His428-Ala581 for GluN2D. ABD S2 was segmented at Gln661-Gln811 for GluN2A, Gln662-Gln812 for GluN2B and Thr686-Lys836 for GluN2D. The C-terminal domain (CTD) for GluN2D was cleaved after Pro876. All chimeric and truncated constructs were generated with a Gibson assembly kit (NEBuilder® M5520AA). All site-direct mutagenesis was introduced by PCR using primers that replaced homologous nucleic acid bases with mutant codons. cRNAs of all these constructs were produced based on restriction enzyme linearized cDNA templates using an RNA transcription kit (Invitrogen® AM1345), and the quality was assessed by gel electrophoresis.

*Two-electrode voltage-clamp recording*

For two-electrode voltage-clamp (TEVC) recording, *Xenopus laevis* oocytes were prepared, injected and voltage clamped as reported (4). In brief, oocytes were injected with 36.8 nL of cRNAs or cDNAs of mixed GluN1-1a and GluN2 subunits at a concentration of 10-50 ng/µL at a 1:1 GluN1:GluN2 ratio. After injection, the oocytes were transferred to Barth solution at 15 °C containing (in mM) 88 NaCl, 10 HEPES, 1 KCl, 2.4 $NaHCO_3$, 0.33 $Ca(NO_3)_2$, 0.41 $CaCl_2$, and 0.82 $MgSO_4$ (pH adjusted to 7.6 with NaOH). Two-electrode voltage-clamp recording was performed 24-48 h after injection and conducted in Ringer solution containing (in mM): 100 NaCl, 0.3 $BaCl_2$, 5 HEPES, and 0.01 DTPA (pH adjusted to 7.3 with KOH with a final potassium concentration of approximately 2.5 mM). The solution was immediately perfused by gravity flow into the animal poles of the oocytes, and the different solutions were exchanged manually towards an 8-modular valve positioner. Both glass electrodes were filled with 3 M KCl solution, and the current responses were recorded using an OC-725 amplifier (Warner Instruments, Hamden, CT, USA) matched with a digital analogue converter (Molecular Devices, Axon™ Digidata® 1550B) under a holding potential of –60 mV unless performing the current–voltage ramp pulse. Glycine (100 µM) and glutamate (100 µM) were regarded as saturating agonists used to induce the current of NMDA receptors unless otherwise stated. We perfused YY-23 for 60 seconds, except for the GluN1-2A and GluN1-2B receptors, to adequately detect its effect. The DMSO used to dissolve the YY-23 was supplemented at the same concentration to each solution used in the same trace, but the total amount did not exceed 0.2%. All traces were analysed with Clampfit 10.6 software, and individual traces were fitted using their own data points with Origin 8.0 for presentation. The current–voltage ramp pulses were recorded by clamping the oocytes successively at 8 different voltages from −100 mV to +40 mV (20 mV apart) within 1 s for each voltage after perfusing the cells with YY-23 for 60 s without withdrawing the drug.

*Patch-clamp recording of HEK 293T cells*

HEK 293T cells (ATCC, Cat. No. CRL-3216) were maintained in DMEM/F12 (Gibco) supplemented with 10% foetal bovine serum (Gibco) and 10 U $ml^{-1}$ penicillin and penicillin–streptomycin (Cellgro) in a humidified incubator at 37 °C (5% $CO_2$). cDNAs encoding the N-methyl-D aspartate receptor (NMDAR) subunits and green fluorescent protein (GFP) at a ratio of 1:1:1 (GluN1:GluN2:GFP) were transiently transfected into HEK293T cells using Fugene HD transfection reagent (Invitrogen) and incubated for 12-16 h. Whole-cell patch-clamp recordings were performed at −60 mV using Axopatch 700B (Molecular Devices). The amplification and data acquisition were performed by an Axon Digidata 1550B (Molecular Devices). The electrodes were filled with an internal solution containing the following (in mM): 110 D-gluconate, 110 CsOH, 30 CsCl, 5 HEPES, 4 NaCl, 0.5 $CaCl_2$, 2 $MgCl_2$, 5 BAPTA, 2 NaATP, and 0.3 NaGTP (pH adjusted to 7.35 with CsOH). The extracellular recording solution was composed of the following (in mM): 150 NaCl, 10 HEPES, 3 KCl, 0.5 $CaCl_2$, and 0.01 EDTA (pH adjusted to 7.4 with NaOH). We used 20 μM glycine and 500 μM NMDA to induce the current of NMDA receptors expressed on HEK 293T cells. The current responses were filtered at 2 kHz (8 pole Bessel filter, −3 dB) and digitized at 20–40 kHz.

*Multiplex fluorescence in situ hybridization*

Mice were perfused intracardially with DEPC-PBS followed by ice-cold 4% PFA in PBS. The brains were dissected, postfixed in 4% PFA for 12-24 hours at 4 °C and dehydrated with 30% sucrose in DEPC-PBS. Afterwards, the brains were sectioned at 20 μm thickness and mounted on SuperFrost Plus® Slides (Thermo Fisher Scientific). Multiplex FISH was then performed with an RNAscope Multiplex Fluorescent v2-Kit (ACDBio). Probes against *grin2d* mRNA, *Slc32a1* mRNA, and *Slc17a7* mRNA were ordered from ACDBio and used in the experiment. Images were captured under a ×20 or ×63 objective using a confocal microscope (Lecia TCS SPS CFSMP).

*Slice preparation*

Mice were anaesthetized with isoflurane, sacrificed and then perfused with 20 ml of ice-cold, oxygenated artificial cerebrospinal fluid (ACSF, 25.0 $NaHCO_3$, 1.25 $NaH_2PO_4$, 2.5 KCl, 0.5 $CaCl_2$, 7.0 $MgCl_2$, 25.0 glucose, 11.0 choline chloride, 11.6 ascorbic acid and 3.1 pyruvic acid in mM, gassed with 95% $O_2$ and 5% $CO_2$). The brains were immediately dissected out after decapitation. Bilateral coronal slices containing the medial PFC (350 μm thick) were cut using a vibratome (Leica VT1000s, USA). The slices were quickly placed into a chamber and incubated in normal oxygenated ACSF (118 NaCl, 2.5 KCl, 26 $NaHCO_3$, 1 $NaH_2PO_4$, 10 glucose, 1.3

$MgCl_2$ and 2.5 $CaCl_2$ in mM, gassed with 95% $O_2$ and 5% $CO_2$) at 35 °C for 2 h. The slices were finally transferred to the recording chamber at room temperature.

*Novelty-suppressed feeding test in mice*

Mice were deprived of food for 24 hours with access to water *ad libitum* before the test was carried out. The mice were then removed to the test room and placed into a clean cage to habituate for at least 1 hour. The testing apparatus consisted of a Plexiglas box (40×40×20 cm), the floor of which was covered with 2 cm of wooden bedding. A small piece of mouse chow was placed on a white platform (white circular filter paper, 11 cm in diameter) positioned in the centre of the box. Each mouse was then placed in a corner of the box with his head facing the platform. The latency of first feeding episode was recorded. The first successful feeding was defined by the mouse sitting on its haunches, holding the pellet with its forepaws and biting the pellets. Immediately after the first feeding, each mouse was transferred to his home cage with access to food *ad libitum,* and the amount of food consumed was measured to control for a change in appetite as a possible confounding factor.

*Force swimming test*

This test was conducted under normal light, as previously described (5). Briefly, mice were individually placed in a cylinder (35 cm height ×10 cm diameter) filled with water. An appropriate water level was set to prevent each mouse from touching the bottom with its limbs. In addition, the water temperature was set at 23-25 °C. The mice were allowed to swim for 6 minutes, and their activity was videotaped. There had been an extra acclimation period with a

10-min swim on the previous day. The duration of immobility, defined as the time spent floating or remaining motionless, was quantified manually during the last 4 minutes.

*Open field test*

The open field *test* is a standard method to profile locomotor activity. Briefly, mice were handled in advance and then placed into the centre of a square box (100 x 100 x 45 cm) that was made of white acrylic plastic. Each mouse was allowed to explore in the box for 10 min and was monitored online. The distance moved and percent of distance travelled in the centre area for each animal were calculated offline.

*Elevated plus maze test in rats and mice*

Mice with acute administration performed this test. The plus maze apparatuses were made of black Plexiglas and had two opposite open arms and two enclosed arms of the same size in a cross-shaped form and in a central region. The four arms of both apparatuses were 70 cm above the ground. The size of the four arms was 27 cm long × 5 cm wide with central square (5 × 5 cm) for mice. The walls of two enclosed arms were at height of 20. Each animal was individually placed on the center platform facing an open arm to initiate the test session. Each animal was allowed to perform for 6 minutes, and the activity was videotaped. Behaviors scored were calculated by percent of the number of entries into open and closed arms. Arm entries were defined as entry of all four paws into the arm.

*Tail suspension test*

The mice were suspended by securing their tails on the edge of a shelf positioned at a height of 80 cm above the floor. Their activity was videotaped to analyse their escape behaviour. A blinded experiment was conducted to record the amount of time spent immobile during the last 4 min of the testing period; the first 2 min were excluded.

*Procedure for the development of a mouse depressive model of chronic social defeat stress (CSDS)*

As previously described (6), in brief, after screening to identify aggressive CD-1 mice, intruder C57BL/6N mice were directly placed within the resident aggressor's home cage compartment for 5-10 min on 10 consecutive days. During this exposure, the intruders showed signs of stress and subordination, including vocalization, the flight response and a submissive posture. After 10 min of social defeat, the intruder was transported across the perforated divider to the opposite compartment and house in this compartment for the remainder of the 24-h period. We placed the control C57BL/6N mice that did not experience social defeat in the same compartment. Twenty-four hours after the last session, we screened the susceptible mice based on the social interaction (SI) ratio. Briefly, SI was obtained by dividing the time spent in the interaction zone when the target was present by the time spent in the interaction zone when the target was absent. An SI ratio of less than 1 indicates that less time was spent in the presence than in the absence of a social aggressor. This can be used a threshold for identifying susceptible mice. All mice were then housed individually for 3 weeks, and all of the susceptible mice were randomly divided into three groups for drug administration. The time spent in the interaction zone was accessed weekly using the same method as previously described (6).

*Electrophysiology in mPFC slices*

Electrophysiological recordings of features including action potentials (APs), sIPSC, mIPSC, fEPSPs and mEPSCs were carried out using an experimental device composed of an Axopatch 700B amplifier (Molecular Devices) and an Olympus microscope (Olympus, Japan) equipped with infrared differential interference contrast optics. The expression of EGFP was visualized in interneurons in the acute slice preparation of GAD67-EGFP knock-in mice using epifluorescence illumination. The APs of interneurons in layer II/III and the mEPSCs of pyramidal neurons in layer V in the prelimbic mPFC were recorded by the whole-cell patch-clamp technique. The glass electrodes had a resistance tip in the range of 3-4 MΩ. For AP recording, the pipettes were filled with intracellular solution containing (in mM) 114 potassium gluconate, 6 KCl, 0.5 $CaCl_2$, 0.2 EGTA, 4 ATP-Mg, and 10 HEPES (pH adjusted to 7.25 with NaOH). AP firing was evoked in current-clamp mode by current injection. For the IPSC recording, the internal solution containing the following (in mM): 130 Cs-methanesulfonate, 10 CsCl, 10 HEPES, 4 NaCl, 7 phosphocreatine, 0.3 Na-GTP, 4 Mg-ATP, and 2 QX314-Br (pH ~7.3 adjust with CsOH). sIPSC were recorded in the mPFC pyramidal neurons voltage clamped at + 10 mV and mIPSC were recorded in the presence of TTX. For the mEPSC recording, neurons were clamped at -65 mV in the presence of TTX (1 μM) and bicuculline (10 μM) in ACSF solution. The intracellular solution contained (in mM) 115 $CsMeSO_3$, 20 CsCl, 10 HEPES, 2.5 $MgCl_2$, 4 Na2-ATP, 0.4 Na-GTP, 10 Na-phosphocreatine, and 0.6 EGTA; pH adjusted to 7.0 with NaOH). The data were filtered at 1 kHz and sampled at 10 kHz using Axon Digidata 1550B (Molecular Devices, USA) and analysed by the Mini Analysis Program (Synaptosoft). For fEPSP recording, the bipolar stimulating electrode was placed on layer V of the mPFC slice, and the recording electrode (filled with 0.5 M $CH_3COONa$, 1-5 MΩ resistance at the tip) was placed on layer II/III. The duration of the stimulus pulse was 100 μs with delivery at 0.03 Hz, and the stimulus intensity was adjusted to 65% of the maximal response. When the evoked responses were maintained at a stable level at this intensity, additional baseline recording was performed for 10 minutes prior to the drug treatment. All recordings were continued for 60-80 minutes. NMDA-evoked currents were recorded in the presence of 200 μM NMDA and 30 μM glycine, which were pressurized and applied in a brief pulse. After 3-5 stable NMDA-evoked currents in the ACSF solution, 3 μM CIQ, 20 μM YY-23 or both were bath-applied for 15-20 min. Current responses were then evoked again by pressure application of NMDA and glycine and compared with current responses obtained during application of the control (ACSF).

*Drug administration*

*For the in vivo* study, freshly prepared YY-23 was suspended in 1% carboxymethyl cellulose sodium (CMC-Na). For acute administration, 1 hour before the behavioural tests, YY-23 was intragastrically (i.g.) administered, and midazolam or fluoxetine (Santa Cruz Biotechnology, USA) was intraperitoneally (i.p.) administered. YY-23 (i.g.), fluoxetine (i.p.) and vehicle (i.g.) were chronically administered (once a day at 9:00 am for 3 weeks) to control and susceptible mice. For the *in vitro* study, YY-23 was first dissolved in dimethyl sulfoxide (DMSO) and then diluted into extracellular recording solution for patch-clamp recording in HEK 293T cells, Ringer solution for TEVC recording or normal ACSF for slice recording. Fluoxetine was dissolved in 0.9% NaCl aqueous solution.

*Protein extraction and filter-aided sample preparation*

Twenty-four hours after the last session of behavioural tests with the mouse depressive model of CSDS, mice were anaesthetized and decapitated immediately. The PFC brain regions were obtained to analyse protein expression by employing quantitative proteomics. The brain tissues were lysed in lysis buffer (4% SDS (m/v), 100 mM DTT, 100 mM Tris, pH = 7.6) and homogenized *via* sonication. The lysate was centrifuged at 14,000 ×g for 20 min, and the supernatant was collected. The protein concentration was determined by tryptophan fluorescence emission assay as described previously (7). Afterwards, approximately 100 µg of protein from each sample was processed by filter-aided sample preparation (FASP) digestion as previously described (8). The tryptic peptides were collected by centrifugation and desalted *via* StageTipC18 purification.

*Nanoflow Liquid Chromatography Tandem Mass Spectrometry*

Proteogenomic analysis of the PFC samples was performed on a Q-Exactive mass spectrometer with an ancillary EASY-nLC 1000 HPLC system (Thermo Fisher Scientific). The tryptic digested peptides were loaded on a 75 µm × 200 mm fused silica column packed in-house with 3 µm ReproSil-Pur C18 beads (Dr. Maisch GmbH, Ammerbuch, Germany) and separated with a 240-min gradient at a flow rate of 250 nL/min. Solvent A was water containing 0.1% formic acid; solvent B was acetonitrile containing 0.1% formic acid. The gradient was 1-4% B, 4 min; 25% B, 210 min; 35% B, 15 min; 90% B, 3 min; and 90% B, 8 min. The mass spectrometry instrument parameters were as follows: MS1 full-scan resolution, 70000 at m/z 200; automatic gain control target, $1 \times 10^6$; maximum injection time, 30 ms; MS2 scan resolution, 17500 at m/z 200; automatic gain control target, $1 \times 10^5$; maximum injection time, 60 ms; isolation window, 2.0 m/z; and dynamic exclusion, 25 s. The precursor ions were fragmented by higher-energy collisional dissociation (HCD) with a normalized collision energy of 30%. NAC samples were analysed on an Orbitrap Elite mass spectrometer with an ancillary EASY-nLC 1000 HPLC system (Thermo Fisher Scientific). MS1 full-scan range, 350.0-1600.0; isolation window, 2.0 m/z; normalized collision energy, 35%.

*Data Analysis*

We calculated the relative current after YY-23 application to evaluate the effects of YY-23 on all wild-type, truncated and other mutant NMDA receptors expressed in recombinant systems. We used $IC_{50}$, the concentration that inhibits the response half-maximally, to evaluate the YY-23 efficiency. The $IC_{50}$ values were determined by nonlinear least-squares fitting of a Hill equation

$$Y = A + \frac{1 - A}{1 + 10^{(LogIC_{50} - [X])^{n_H})}}$$

with concentration–response data from individual trials normalized to the maximal responses in the absence of inhibitor. In this equation, A is the minimal plateau of the dose–response curve; the relative current (Y) was measured at the 60th second of YY-23 application on oocytes and at the current plateaus for HEK293T cells; X is the YY-23 concentration; and $n_H$ is the Hill slope. Fitted $IC_{50}$ and Hill slope values for individual oocytes were used to calculate the mean and SEM as the final $IC_{50}$ and Hill slope values. For voltage–current data analysis, we present the mean values with their associated SEM error bars.

To screen the pharmacological targets for YY-23 among monoaminergic receptors, calcium flux was measured by the FLIPR signal (FLIPR Calcium 4 Assay Kit), and cAMP was measured by the TR-FRET signal (Lance Ultra cAMP kit). The data are reported as the mean from the experiment. EC50 (log(agonist) *vs.* response - variable slope): Y = bottom + (top-bottom)/(1+10^((logEC50-X) * Hill slope). IC50 (log(inhibitor) *vs.* response - variable slope): Y = bottom + (top-bottom)/(1+10^((logIC50-X) * Hill slope). GraphPad Prism software was used for data analysis and EC50 or IC50 calculation. Activation % = $\frac{\text{raw data} - \text{minimum signal}}{\text{maximum signal} - \text{minimum signal}} \times 100$ and inhibition % = $100 - \frac{\text{raw data} - \text{minimum signal}}{\text{maximum signal} - \text{minimum signal}} \times 100$ for the targets $5HT_{1A}$, $D_2R$, $D_4R$, $5HT_{2A}$, $5HT_{2B}$, and $5HT_{2C}$; activation % = $100 - \frac{\text{raw data} - \text{minimum signal}}{\text{maximum signal} - \text{minimum signal}} \times 100$ and inhibition % = $\frac{\text{raw data} - \text{minimum signal}}{\text{maximum signal} - \text{minimum signal}} \times 100$ for the target $D_1R$.

The MS data were analysed *via MaxQuant* software (http://maxquant.org/, version 1.6.5.0). Carbamidomethyl (C) was set as a fixed modification, while oxidation (M) and protein N-terminal acetylation were set as variable modifications. The MS/MS spectra were searched against the Mouse UniProt FASTA database (UP000000589, 2019, containing 55153 entries). Trypsin/P was selected as the digestive enzyme, with two maximum allowed missed cleavages. The false discovery rate (FDR) for peptides and proteins was controlled at <1% by the Andromeda search engine. A Venn plot was plotted *via* Jvenn (http://jvenn.toulouse.inra.fr/app/example.html). Principal component analysis (PCA) was performed with MetaboAnalyst (https://www.metaboanalyst.ca/MetaboAnalyst/home.xhtml). Functional annotations were performed with the Database for Annotation, Visualization, and Integrated Discovery (DAVID; https://david.ncifcrf.gov/), and the results were visualized *via* R studio. The GO pathways were visualized with Pathview (https://pathview.uncc.edu/guest). Student's *t* test was used to screen out differentially expressed proteins between different groups, and a $p$ value < 0.05 indicated significance. A fold change >1.2 was considered to indicate significantly different expression. The raw proteomics data have been deposited to the ProteomeXchange (9) Consortium *via the* PRIDE (10) partner repository with the dataset identifier.

*Statistical Analyses*

Data meeting the assumptions of normality and equal variance between sample groups are shown as the mean ± SEM. Data that did not meet the normality or equal variance assumptions are presented in box plots with bars showing the medians and boxes showing the lower/upper quartiles. All statistical tests were two-tailed, and significance was assigned at $p < 0.05$. The analyses were blindly performed for treatment assignments in all behavioural experiments. The statistical analyses were performed using GraphPad Prism v6 software. Normality and homogeneity of variance between group samples were assessed using the Shapiro–Wilk test and Bartlett's test, respectively. Once normality and equal variance between sample groups were achieved, *t* tests, one-way and two-way ANOVA followed by *post hoc* multiple comparisons tests were used. When normality tests failed, Mann–Whitney U, Wilcoxon, Kruskal–Wallis one-way ANOVA and Friedman's two-way ANOVA tests were performed. When homogeneity of variance tests failed, unpaired t test with Welch's correction and Brown-Forsythe ANOVA was used. In rare cases in which values were missing randomly from repeated-measures samples, the data were analysed by fitting a mixed-effect model in Prism 9.

### Supplementary figures and figure legends

**Fig. S1**

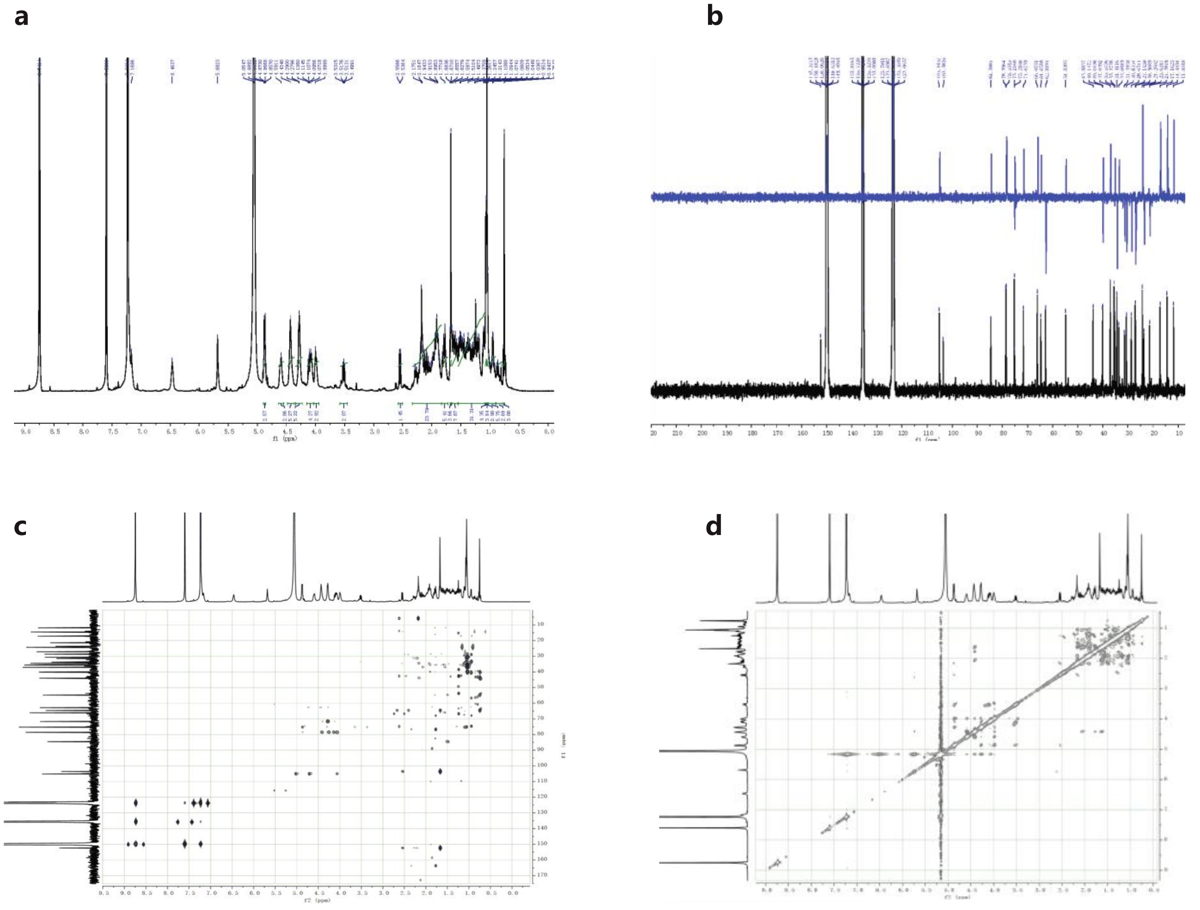

Fig. S1| **Spectroscopic analysis of YY-23.** $^{1}$H (**a**, 400 MHz, in pyridine-$d_5$), $^{13}$C (**b**, 100 MHz, in pyridine-$d_5$) NMR, HMBC (**c**) and ROESY (**d**) spectra of YY-23.

### Fig. S2

**a**

```
                         M1                                        M2
2A  (557) VWVMMFVMLLIVSAIAVFVF      (598) SFTIGKAIWLLWGLV
2B  (558) VWVMMFVMLLIVSAVAVFVF      (599) SFTIGKAIWLLWGLV
2C  (555) VWVMMFVMCLTVVAITVFMF      (596) SFTIGKSVWLLWALV
2D  (582) VWVMMFVMCLTVVAVTVFIF      (623) TFTIGKSIWLLWALV
                   * * *   *                    *      *
                         M3                                        M4
2A  (629) IMVSVWAFFAVIFLASYTANLAAFM      (815) DNMAGVFYMLAAAMALSLITFIWEHL
2B  (630) IMVSVWAFFAVIFLASYTANLAAFM      (816) DNMAGVFYMLGAAMALSLITFICEHL
2C  (627) IMVLVWAFFAVIFLASYTANLAAFM      (813) DNMAGVFYMLLVAMGLALLVFAWEHL
2D  (654) IMVLVWAFFAVIFLASYTANLAAFM      (840) DNMAGVFYMLLVAMGLSLLVFAWEHL
             *                                            **   *    ** *
```

**b**

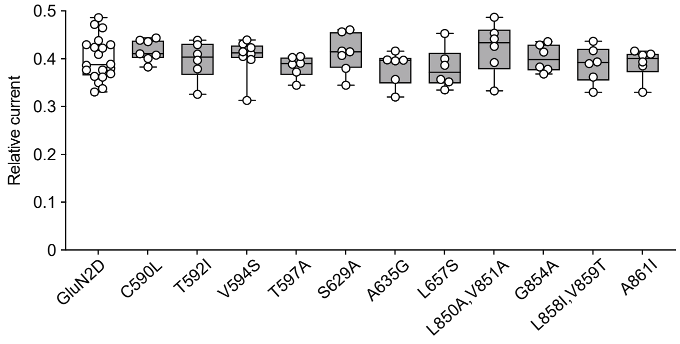


Fig. S2| **The transmembrane domain of the GluN2D subunit is not responsible for the inhibition of YY-23.** (**a**) Sequence alignment of the four transmembrane helices of the rat GluN2 subunits (2A to 2D). Nonconserved residues between GluN2A (2B) and GluN2C (2D) are marked by asterisks under the amino acid columns. (**b**) Inhibitory effect of 10 μM YY-23 on the saturating agonist-induced activity of NMDA receptors containing wild-type GluN1 subunits and GluN2D subunits with various mutations. The current activity with saturating agonists was normalized to 1. There were no significant differences between wild-type and individual mutant receptors. The data were analysed by Kruskal–Wallis one-way ANOVA.

**Fig. S3**

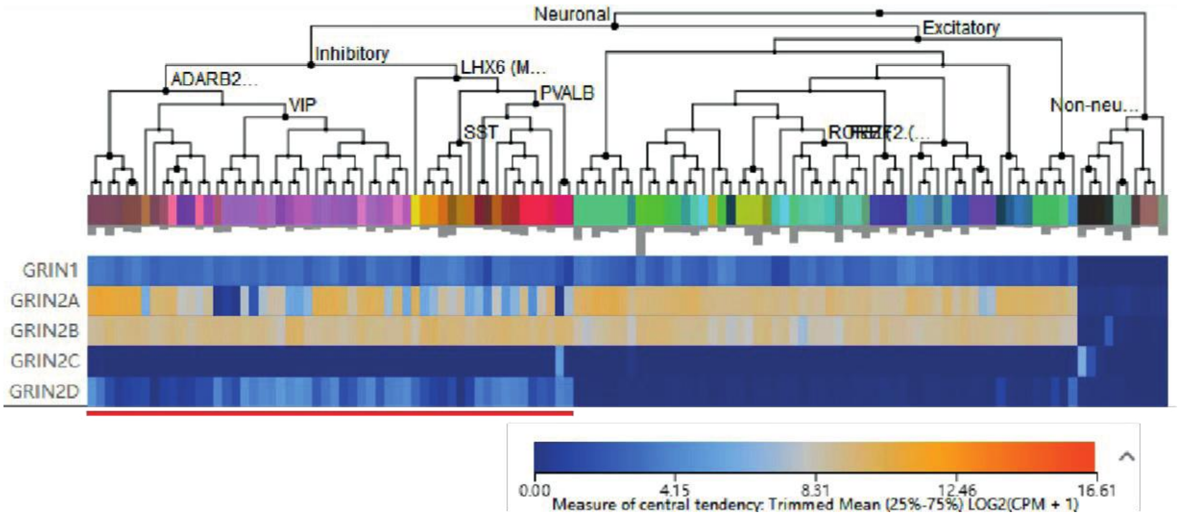


Fig. S3| **Major NMDAR subunits mRNA distribution in different neuronal cell type**. Data was derived from a dataset of mouse whole cortex and hippocampus 10X genomics RNAseq from Allen Brain Atlas. mRNA expression level of five major subunits of NMDARs were analyzed using single cell RNA-seq database. Data was analyzed and presented online with 10X-SMART-SEQ TAXONOMY (https://portal.brain-map.org/atlases-and-data/rnaseq).

**Fig.S4**

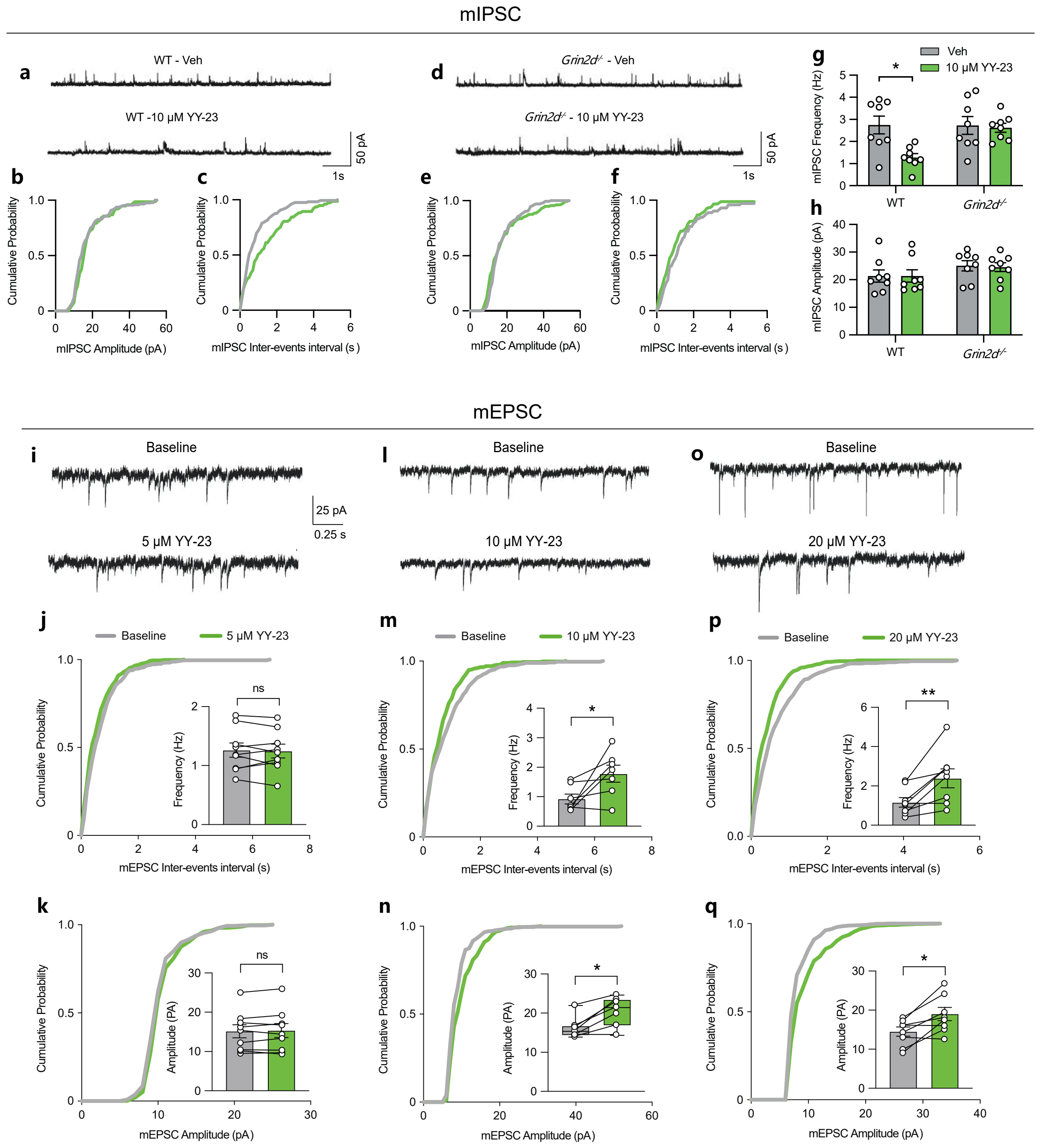


Fig. S4| **YY-23 reduced mIPSC and increased the mPFC pyramidal neuron activity**. **(a, d)** Example traces showing mIPSC recorded from pyramidal neurons in the WT and KO mice. **(b, c, e, f)** The cumulative distribution of the mIPSC amplitude and interevent intervals. The data were analysed by Kolmogorov–Smirnov test. **(g, h)** The bar graphs represent the average frequency

(Hz) and amplitude (pA) after vehicle (grey bars) and after YY-23 treatment (green bars). **(i, l, o)** Example of mEPSC traces shown in the presence of 5 μM, 10 μM or 20 μM YY-23 after 5 minutes of baseline recording. The data were measured in a whole-cell configuration from the mPFC pyramidal neurons. The cumulative distribution of the mEPSC the interevent intervals **(j, m, p)** and amplitude **(k, n, q)** are shown. The data were analysed by Kolmogorov–Smirnov test. The inseted bar graphs represent the average frequency and amplitude at baseline (grey bars) and after vehicle and YY-23 treatments (green bars). n.s. means no significant difference between baseline and treatment. The bar graphs show the mean ± SEM, and the data were analysed by paired t test. The box plot graph shows the lower/upper quartiles, and the lines represent the median values; the data were analysed by Wilcoxon tests. *$p < 0.05$, and **$p < 0.01$.

### Fig. S5

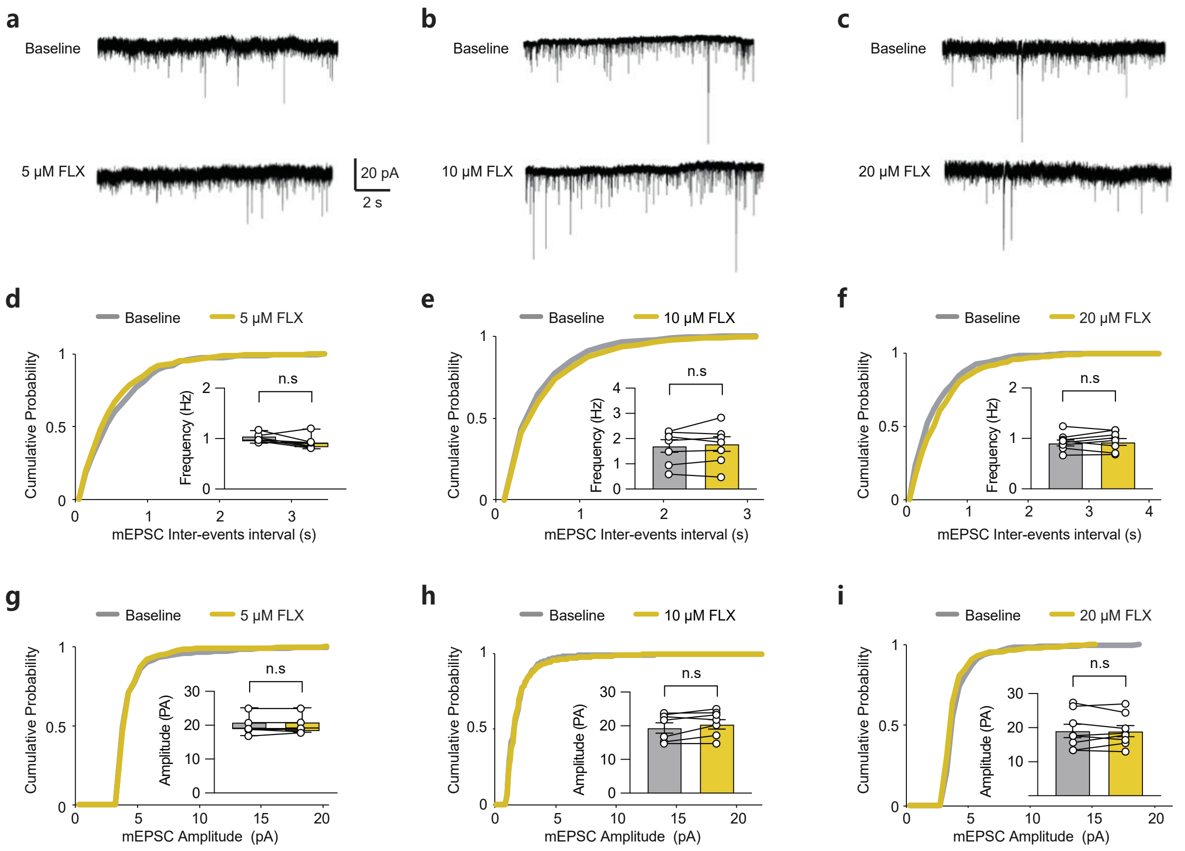


Fig. S5| **Fluoxetine did not change mPFC pyramidal neuron activity.** (**a-c**) Example of mEPSC traces shown in the presence of 5 µM, 10 µM or 20 µM fluoxetine after 5 minutes of baseline recording. The data were measured in a whole-cell configuration from the mPFC pyramidal neurons. The cumulative distribution of the mEPSC amplitude **(d-f)** and the interevent intervals **(g-i)** are shown. The data were analysed by Kolmogorov–Smirnov test. The inset bar graphs represent the average frequency (Hz) and amplitude at baseline (grey bars) and after fluoxetine treatment (yellow bars). n.s. means no significant difference between baseline and treatment. The bar graphs show the mean ± SEM, and the data were analysed by paired t test. In the box plots, the bars show the min/max values, the boxes show the lower/upper quartiles, and the lines represent the median values. The data were analysed by Wilcoxon test.

**Fig. S6**

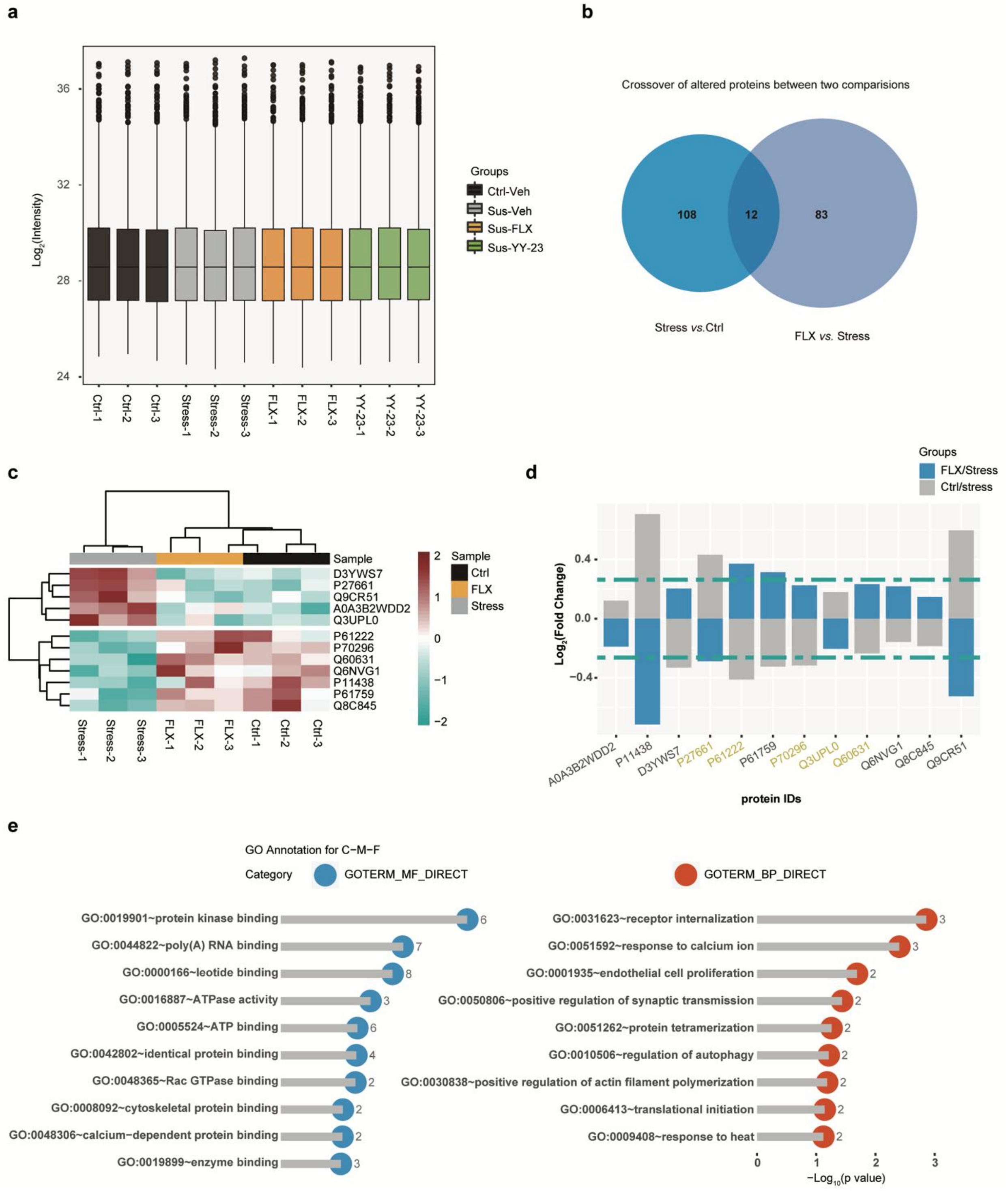


Fig. S6| **Proteomic analysis of fluoxetine administration.** (**a**) Distribution diagram showing the variations in protein expression among the four groups and indicating that there was no significant variation in total protein expression among the groups. (**b**) Venn diagram revealing the numbers of altered proteins in the Sus-Veh/Ctrl-Veh comparison and the Sus-FLX/Sus-Veh comparison. Note that the administration of fluoxetine in susceptible mice rescued the levels of only 12 proteins. (**c**) Heatmap clustering of 12 proteins showing that the levels of the altered proteins in the susceptible mice (Stress 1-3) were rescued by the administration of fluoxetine. (**d**) Fold changes in protein expression between the Sus-Veh and Ctrl-Veh groups and the Sus-FLX and Sus-Veh groups. (**e**) GO enrichment analysis showed the molecular functions and biological processes associated with the proteins shown in (**c**).

### Fig. S7

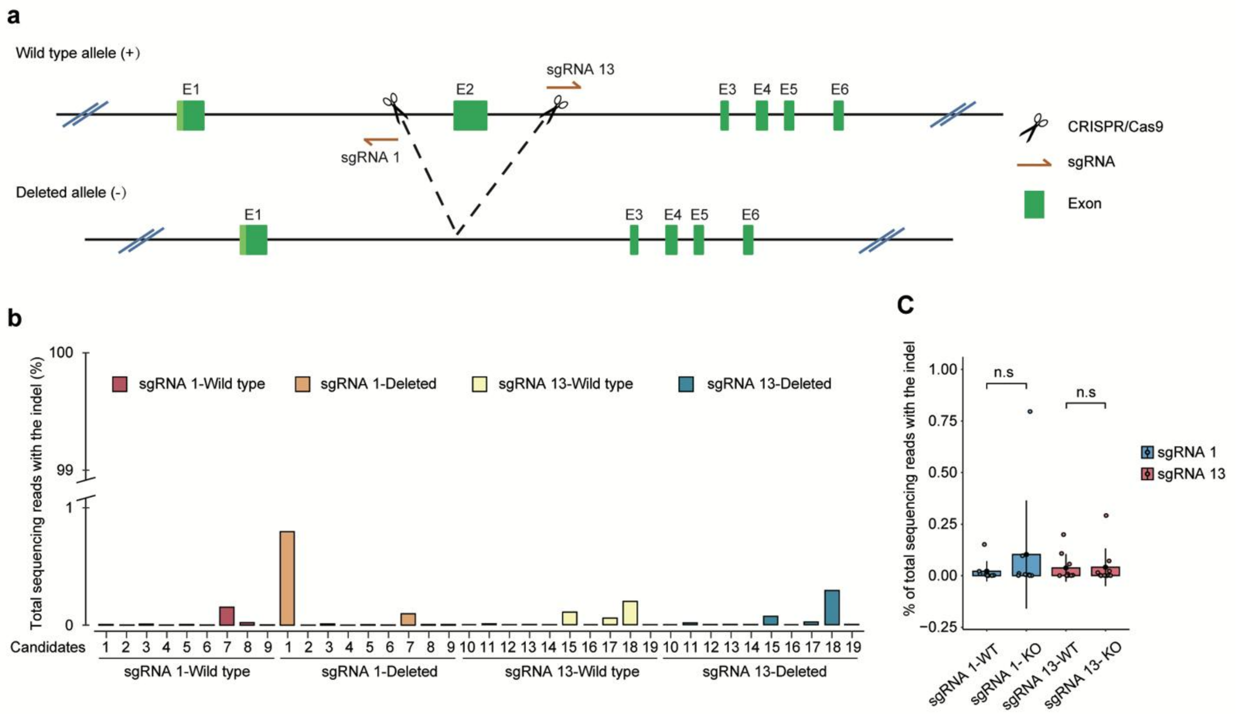


Fig. S7| **Transgenic mouse construction and validation**. (**a**) Schematic diagram of the *grin2d*-knockout mouse design strategy. Exon 2 of *grin2d* was deleted with the CRISPR/CAS9 system guided by two sgRNAs. (**b**) Editing outcomes of 19 candidates targeting different sgRNAs (numbers 1-9 for sgRNA 1 and 10-19 for sgRNA 13) in WT and *grin2d*-KO mice. (c) Comparison of off-target effects between WT and KO mice. The *n.s* shown above the columns indicates that there was no difference in the indel probability between KO and WT mice with different sgRNAs (n = 9 for sgRNA1 and n=10 for sgRNA13). The data were analysed by Friedman's two-way ANOVA. In the box plots, the bars show the min/max values, and the boxes show the lower/upper quartiles.

Table S1. The [1]H (400 MHz) and [13]C (100 MHz) NMR spectroscopic data for YY-23 (pyridine-$d_5$)

| Position | $\delta_C$ (ppm) | $\delta_H$ (ppm) | Position | $\delta_C$ (ppm) | $\delta_H$ (ppm) |
|---|---|---|---|---|---|
| 1 | 30.6 | 1.89 (m) | 18 | 14.8 | 0.71 (s) |
| 2 | 27.1 | 1.94 (m),1.33 (m) | 19 | 24.4 | 0.85 (s) |
| 3 | 66.0 | 4.42 (m) | 20 | 104.1 | - |
| 4 | 34.4 | 1.63 (m),1.55 (m) | 21 | 12.3 | 1.64 (s) |
| 5 | 37.0 | 2.17 (m) | 22 | 152.9 | - |
| 6 | 26.9 | 1.34 (m),1.18 (m) | 23 | 31.1 | 2.07 (m), 1.56 (m) |
| 7 | 28.6 | 1.76 (m),1.59 (m) | 24 | 24.1 | 2.26 (m), 1.04 (o) |
| 8 | 35.2 | 1.52 (m) | 25 | 34.2 | 1.97 (m) |
| 9 | 40.2 | 1.80 (m) | 26 | 75.8 | 4.05 (o), 3.50 (m) |
| 10 | 35.7 | - | 27 | 17.7 | 1.04 (d, J=6.8 Hz) |
| 11 | 21.8 | 1.36 (m), 1.21 (o) | Glu | | |
| 12 | 40.6 | 1.76 (m), 1.20 (o) | 1′ | 105.1 | 4.87 (m, J=7.7 Hz) |
| 13 | 44.4 | - | 2′ | 75.2 | 4.07 (m) |
| 14 | 55.2 | 0.88 (o) | 3′ | 78.5 | 4.28 (m) |
| 15 | 31.9 | 1.87 (m), 1.49 (m) | 4′ | 71.6 | 4.28 (m) |
| 16 | 85.1 | 4.85 (o) | 5′ | 78.6 | 4.00 (m) |
| 17 | 65.2 | 2.51 (m) | 6′ | 62.8 | 4.59 (m),4.43 (m) |

Table S2. Screening pharmacological target for YY-23 among monoaminergic receptors

| Compound name | Assay name | Assay target | Bottom | Top | LogEC 50 | $EC_{50}$ (M) |
|---|---|---|---|---|---|---|
| 8-OH-DPAT | Agonist | 5-HT1A | 9.3 | 92.9 | -8.7 | 2.04E-09 |
| a-methyl-5HT | Agonist | 5-HT2A | 2.1 | 121.5 | -7.4 | 3.62E-08 |
| a-methyl-5HT | Agonist | 5-HT2B | -17.5 | 117.0 | -7.5 | 3.29E-08 |
| a-methyl-5HT | Agonist | 5-HT2C | -6.2 | 105.5 | -7.7 | 2.14E-08 |
| Dopamine | Agonist | D1 | 0.1 | 99.4 | -8.6 | 2.74E-09 |
| Dopamine | Agonist | D2 | 1.4 | 102.6 | -8.2 | 6.77E-09 |
| Dopamine | Agonist | D4 | 1.6 | 102.8 | -7.6 | 2.29E-08 |
| YY-23 | Agonist | 5-HT1A | 0.0 | 0.0 | 0.0 | no fit |
| YY-23 | Agonist | 5-HT2A | 0.0 | 0.0 | 0.0 | no fit |
| YY-23 | Agonist | 5-HT2B | 0.0 | 0.0 | 0.0 | no fit |
| YY-23 | Agonist | 5-HT2C | 0.0 | 0.0 | 0.0 | no fit |
| YY-23 | Agonist | D1 | 0.0 | 0.0 | 0.0 | no fit |
| YY-23 | Agonist | D2 | 0.0 | 0.0 | 0.0 | no fit |
| YY-23 | Agonist | D4 | 0.0 | 0.0 | 0.0 | no fit |
| WAY100135 | Antagonist | 5-HT1A | 5.2 | 108.5 | -6.4 | 3.76E-07 |
| Risperidone | Antagonist | 5-HT2A | 19.8 | 98.2 | -8.8 | 1.72E-09 |
| Methiothepin | Antagonist | 5-HT2B | 12.1 | 102.2 | -7.0 | 1.08E-07 |
| Risperidone | Antagonist | 5-HT2C | -4.3 | 99.4 | -8.3 | 4.80E-09 |
| SCH-23390 | Antagonist | D1 | 3.7 | 102.8 | -9.0 | 9.48E-10 |
| Risperidone | Antagonist | D2 | -16.2 | 96.8 | -8.8 | 1.61E-09 |
| Clozapine | Antagonist | D4 | -3.4 | 111.3 | -6.4 | 4.14E-07 |
| YY-23 | Antagonist | 5-HT1A | 0.0 | 0.0 | 0.0 | no fit |
| YY-23 | Antagonist | 5-HT2A | 0.0 | 0.0 | 0.0 | no fit |
| YY-23 | Antagonist | 5-HT2B | 12.8 | = 100.0 | -4.9 | > 10uM |
| YY-23 | Antagonist | 5-HT2C | 0.0 | 0.0 | 0.0 | no fit |
| YY-23 | Antagonist | D1 | 0.0 | 0.0 | 0.0 | no fit |
| YY-23 | Antagonist | D2 | 0.0 | 0.0 | 0.0 | no fit |
| YY-23 | Antagonist | D4 | 0.0 | 0.0 | 0.0 | no fit |

Table. S3 $IC_{50}$ values for YY-23 at NMDA receptors expressed on *Xenopus* oocytes and HEK293T cells.

| NMDA receptor subtype | Cell type | $IC_{50}$±SEM | $n_H$ | *N* |
|---|---|---|---|---|
| GluN1-GluN2C | *Xenopus* oocytes | 3.06±0.83 | -1.21 | 4 |
| GluN1-GluN2D | *Xenopus* oocytes | 3.73±0.27 | -2.23 | 4 |
| GluN1-GluN2C | HEK293 cells | 9.30±3.24 | 0.92115 | 5 |
| GluN1-GluN2D | HEK293 cells | 5.22±2.65 | -4.90 | 6 |
